# Population structure and environmental niches of *Rimicaris* shrimps from the Mid Atlantic Ridge

**DOI:** 10.1101/2021.06.27.450074

**Authors:** Pierre Methou, Ivan Hernández-Ávila, Cécile Cathalot, Marie-Anne Cambon-Bonavita, Florence Pradillon

**Author notes:** Current address: X-STAR, Japan Agency for Marine-Earth Science and Technology (JAMSTEC), Yokosuka, 237-0061, Japan. Current address: Facultad de Ciencias Naturales, Universidad Autónoma del Carmen, 14158, Cuidad del Carmen, México (Ivan Hernández-Ávila).

## Abstract

Among the endemic and specialized fauna from hydrothermal vents, *Rimicaris* shrimp surely constitutes one of the most important and emblematic components of these ecosystems. In the Mid Atlantic Ridge, two species affiliated to this genus co-occur: *Rimicaris exoculata* and *Rimicaris chacei* that differ by their morphology, their trophic regime as well as by their abundance. The first forms large and dense aggregations on active vent chimney walls in close proximity to vent fluid emissions, whereas the second is recognized as much less conspicuous, living mostly in scattered groups or solitary further away from the fluids. However, the recent revision of *Rimicaris* juvenile stages from the Mid Atlantic Ridge shows that *R. chacei* abundance would be higher than expected at these early life stages. Here, we describe and compare the population structures of *R. exoculata* and *R. chacei* at the Snake Pit and TAG vent fields. We show distinct population demographics between the two co-occurring shrimps with a large post settlement collapse in *R. chacei* population suggesting a large juvenile mortality for this species. We also observe important spatial segregation patterns between the two species and their different life stages. Additionally, our results highlight distinct niches for the earliest juvenile stages of both *R. exoculata* and *R. chacei*, compared to all the other life stages. Finally, we discuss the potential factors - predation and competitive interactions among others - that could explain the differences we observe in the population structure of these two species.

## Introduction

Fueled by the chemosynthetic energy arising from the mixing between vent fluids and surrounding seawater, hydrothermal vent ecosystems host dense and lush communities of endemic fauna which are organized in distinct species assemblages along steep thermal and chemical gradients. Distributed patchily along mid oceanic ridges, these environments constitute isolated but also ephemeral oases of life within the deep sea, either because of catastrophic events leading to the collapse of entire populations (Fornari *et al*., 2012) or simply because of the general lifespan of a vent field, as fluid activity is doomed to ultimately wane and cease over geological timescales (Van Dover, 2019). Therefore, vent species survival over successive generations is greatly interlinked with their ability to successfully settle in and colonize new vent fields, implying adaptations at each stage of their lifecycle.

As in other marine benthic ecosystems (Menge, 1991), population demographics of species from hydrothermal vents depend on the number of competent larvae reaching the vent site, the probability of success of their settlement and any mortality events occurring between their successive life stages (Kelly and Metaxas, 2007). Therefore, any biotic or abiotic factor affecting one of these three steps will likely impact the capacity of individuals to reach sexual maturity and the size of the adult populations. These environmental factors are acting differentially depending on the life stages considered as each of them can occupy its own ecological niche and differs by its vulnerability to the different mortality factors (Werner and Gilliam, 1984).

Whereas a high juvenile mortality, mostly related to size selective predation, is often characteristic of the post settlement phase in shallow water species (Gosselin and Qian, 1997), a similar generalization has not yet been established in hydrothermal vent ecosystems. This can be attributed to the difficulty of conducting long term population survey and temporal observations along appropriate timescales in such remote places. Settlement and recruitment patterns of these species have then mostly been inferred from cohort monitoring. According to these studies, populations of some vent species such as *Bathymodiolus azoricus* mussels or some gastropods are largely dominated by small size individuals (Comtet and Desbruyères, 1998; Kelly and Metaxas, 2007) implying either an important juvenile mortality or large variations between the different recruitment events. Conversely, a limited number of juveniles has been reported for several others including alvinellid polychaetes and many peltospirid or lepetodrilid gastropods (Zal *et al*., 1995; Faure *et al*., 2007; Matabos and Thiebaut, 2010; Marticorena *et al*., 2020) suggesting a frequent arrival of larvae at the vent field followed by a fast growth rate and an accumulation within the adult populations. Still, spatial or temporal limitation in sampling could also bias our understanding of post settlement processes in these populations, as different life stages sometimes inhabit different parts of the field. Important spatial zonation between juveniles and adults has been reported for instance in *Kiwa tyleri* crabs in which adult aggregates locate near the fluid exits while juveniles remain restricted at the periphery of the field (Marsh *et al*., 2015). Similarly, a segregation according to the organism size was observed along the thermal gradient in *Bathymodiolus azoricus* mussels with juveniles and small mussels preferentially found in areas with lower temperatures (Husson *et al*., 2017).

*Rimicaris* shrimps are important components of hydrothermal vent ecosystems as they constitute some of the most visually dominant megafauna in several regions such as in the Central Indian Ridge (Watanabe and Beedessee, 2015), the Mid Cayman Rise (Plouviez *et al*., 2015) or the Mid Atlantic Ridge (Zbinden and Cambon-Bonavita, 2020). In the northern part of the Mid Atlantic Ridge, two species of this genus co-occur: *Rimicaris exoculata*, forming large aggregations close to vent fluid emissions, with densities of several thousands of individuals per m^2^, and *Rimicaris chacei* supposedly less abundant than its congener (several tens to hundreds of individuals per m^2^) as they are less conspicuous within the vent fields (Segonzac, de Saint Laurent and Casanova, 1993).

It is assumed that population demography of congeneric species should be comparable especially when they evolve in similar habitat conditions (Münzbergová, 2013). Observations of contrasting population sizes and distribution patterns in many biologically similar congeners suggest, however, that distinct population dynamics can exist even in cases of syntopy when species inhabit jointly the same habitat at the same time (Münzbergová, 2013; Jones and Ricciardi, 2014; Bouchemousse, Lévêque and Viard, 2016). A stable co-occurrence at a small spatial scale of closely related species usually presupposes some differentiation in resource use as well as mechanisms that prevent interbreeding (Beermann and Franke, 2012). When congeners display similar biological traits and ecological niches, competitive processes are expected to be intensified (Franke, Gutow and Janke, 2007; Rius, Turon and Marshall, 2009), although they do not necessarily constitute a major driver of their population dynamics (Bouchemousse, Lévêque and Viard, 2016). The two *Rimicaris* species differ by their trophic regime, highly dependent on the dense and diversified chemosynthetic episymbiotic communities hosted in their enlarged cephalothorax *for R. exoculata* (Gebruk *et al*., 2000; Ponsard *et al*., 2013; Zbinden and Cambon-Bonavita, 2020) and relying on symbiotrophy, bacterivory and scavenging for *R. chacei* (Gebruk *et al*., 2000; Apremont *et al*., 2018). This mixotrophic regime of *R. chacei* has been hypothesized to result from the competition for space with *R. exoculata* that would maintain *R. chacei* at a distance from vent fluids essential for fueling their symbionts (Apremont *et al*., 2018).

Zonation patterns between *R. exoculata* life stages have been reported by different studies (Segonzac, de Saint Laurent and Casanova, 1993; Shank, Lutz and Vrijenhoek, 1998; Gebruk *et al*., 2010) which highlighted the existence of red juvenile patches adjacent to the dense aggregations of adults in several MAR vent fields. More recently, spatial segregation between males, found at the periphery away from the fluid mixing zones, and females, mainly occurring within the dense aggregates on active chimney walls, have also been observed (Hernández-Ávila *et al*., 2021). In addition, dense aggregations of small juveniles were also collected close to peripheral and diffuse fluid emissions arising from cracks at the TAG vent field (Methou *et al*., 2020; Hernández-Ávila *et al*., 2021). Surprisingly, these individuals were revealed to be juvenile stages of *R. chacei* unveiling a misidentification in the previous species affiliation of these life stages (Methou *et al*., 2020; Hernández-Ávila *et al*., 2021). Such a large abundance of *R. chacei* early life stages was unexpected considering the low abundance of the adults, and implied for *R. chacei* either a low post-settlement survival, or a larger adult population than previously stated, or both. Therefore, a full characterization of the population structures of *R. exoculata* and *R. chacei* is required to properly assess the demography of each species and whether the differences in the relative abundance of life stages could be related to their different trophic strategies or to other external factors such as predation, or physiological stress arising from their environment.

In this study, we describe and compare the population structures of *R. exoculata* and *R. chacei* from different types of shrimp assemblages at two mid-Atlantic vent fields, TAG and Snake Pit, using length-frequency analysis and the recently revised identification of *Rimicaris* shrimp life stages from these areas (Methou *et al*., 2020). Taking into account the significant spatial zonation of the different life stages of these two species, we also characterized their realized environmental niches and discussed the role of potential biotic and abiotic factors affecting the demography of these shrimp populations.

## Materials & Methods

### Field Sampling

*Rimicaris* shrimps were collected on board the R/V *Pourquoi pas?* during the BICOSE 2 cruise (http://dx.doi.org/10.17600/18000004) between January 26^th^ and March 10^th^, 2018 at the TAG (26°08.2’N, 44°49.5’W, 3620 m depth) and the Snake Pit (23°22.1’N 44°57.1’W, 3470 m depth) hydrothermal vent fields using the suction sampler of the HOV *Nautile*. Samples were collected all over the active venting areas at the TAG vent field and on the three active edifices of Snake Pit: the Beehive, the Moose and the Nail (Figure S1). In total, 3919 *R. exoculata* shrimps (1533 from TAG and 2386 from Snake Pit) and 1201 *R. chacei* shrimps (886 from TAG and 315 from Snake Pit) were collected.

Spatially discrete samples were targeted to explore the fine scale variations in the distribution of populations of each species – from 10’s of centimeters for high densities areas to few meters in sparsely populated ones – with 13 different spatial samples at TAG and 17 at Snake Pit (Table S1). These samples were classified according to the type of vent fluid emissions and their proximity to them. During the dives, each sample was kept separate using the different chambers of the suction sampler carousel and the closeable bioboxes of the HOV *Nautile*. All shrimps were stored in 80% EtOH. For 20 of the sampling points, temperature measurements were carried out prior to shrimps sampling, moving the HOV temperature probe across several points (0.1°C precision for a temperature below 50°C) within the targeted sampling area, for 3 to 6 min 45s with a sampling frequency of 1Hz.

Additionally, a more precise environmental characterization of the vent fluid chemistry was obtained for 14 of these sampling points with pH, iron – Fe^2+^ and total iron (Fe_tot_) – and H_2_S concentrations measurements. Water samples were collected with the PEPITO sampler implemented on the HOV *Nautile* (Sarradin *et al*., 2007) and equipped with blood bags (Terumo^®^). The suction inlet was coupled to the HOV temperature probe. Prior to use, all the equipment used for sampling was rigorously washed 3 times with diluted hydrochloric acid (pH 2, Suprapur, Merck) and then thoroughly rinsed with ultrapure water (Mili-Q element system). Immediately after the recovery of the HOV, samples were processed in the chemical lab onboard (clean lab, P 100 000; ISO8) and pH was measured on a subsample using a Metrohm pH-meter. Measurements were carried out after calibration of the pH-meter with NBS buffers (pH 4, 7 and 10) at 25°C. Total iron (Fe_tot_) concentration were determined onboard on water samples taken by PEPITO using the Ferrozine spectrophotometric method (Stookey, 1970). Free inorganic sulfides (ΣS = HS^-^+S^2-^+H_2_S; (Le Bris *et al*., 2000)) were also determined onboard on recovered water samples using the Cline spectrophotometric method (Cline, 1969).

In addition to the PEPITO sampling, total dissolvable Fe^2+^ were measured *in situ* using a chemical miniaturized analyserr (CHEMINI; (Vuillemin *et al*., 2009)). The *in-situ* measurement was based on flow injection analysis with a colorimetric detection (methylene blue method). A calibration of the analyser was performed *in situ* at the beginning and at the end of each HOV dive using Fe(II) stock solutions. Hydrothermal samples were pumped without any filtration and the signal acquisition (∼ 3 min) was initiated at the same time as the water sampling since the same inlet was used for both PEPITO and CHEMINI. Finally, the *in-situ* concentration of dissolved oxygen was monitored during each sampling using an optode (Aanderaa).

### Life stage identifications and measurements

For each individual, carapace length (CL) was determined to the nearest 0.1 mm by Vernier calipers from the rear of the eye socket to the rear of the carapace in the mid-dorsal line. Each shrimp was identified and sorted according to its life stage and sex. Sex was identified in adults by the occurrence of the appendix masculina on the second pleopod in males, and by the shape of the endopod of the first pleopod (Komai and Segonzac, 2008). Since these sexual characters appear only at the adult stage, sex of juvenile/subadult specimens could not be determined.

Juvenile stages of *R. exoculata* and *R. chacei* were identified according to (Methou *et al*., 2020). Because pleopod morphologies are identical between subadults and small females, the distinction between them was made according to the Onset of Sexual Differentiation (OSD) size defined previously for *R. exoculata* and *R. chacei*, respectively for a CL of 10 mm and 5.98 mm (Methou *et al*., 2020; Hernández-Ávila *et al*., 2021). All the individuals without male characteristics were sorted as females if their size exceeded OSD, whereas individuals with a smaller size than OSD were sorted as subadults. Subadults and juvenile stages were sometimes considered together as immature individuals in the analyses.

### Statistical analysis of the population structure

*Rimicaris* shrimps were grouped first by species and then, by life stage for each vent field. Visual examination of our dataset and Shapiro-Wilk normality tests revealed that the size distributions of the shrimps collected during the BICOSE 2 expedition were not following a Gaussian distribution. Therefore, non-parametric tests were used for intergroup comparisons, using a Mann Whitney test to compare differences in size between vent fields (Snake Pit vs TAG), species (*R. exoculata* vs *R. chacei*) or sex (females vs males). Size frequency distributions between groups were compared with the Kolmogorov-Smirnov two-sample goodness of fit tests. Significant variations in the proportions of the two *Rimicaris* species or in proportions of immature over adult individuals for each of these two *Rimicaris* species were tested using χ^2^ test with Yate’s correction for one degree of freedom. Frequencies of males and females in samples were tested for significant variation from a 1:1 sex ratio using χ^2^ test with Yate’s correction for one degree of freedom (Detailed p-values for these tests are given in electronic supplementary material, table S2). The different spatial samples were grouped by k-means hierarchal clustering based on Euclidean distance matrix and displayed with a heatmap. The optimal number of groups was determined both visually and statistically using the gap statistic method (Tibshirani, Walther and Hastie, 2001). All tests were performed using R version 3.6.1 (R Core Team, 2020).

### Cohort analysis of the population structure

Histograms of the size frequency distributions were analyzed as mixtures of normal distributions for samples containing at least 40 individuals. Size-frequency histograms were plotted using a 1 mm size class interval. This interval was chosen to meet the following criteria described by (Jollivet *et al*., 2000): (1) most size-classes have at least five individuals each, (2) adjacent empty size-classes have been minimized and (3) an interval clearly larger than the estimated error rate on measurements. We determined the overlapping component distribution that gives the best fit to the histogram with the “mixdist” R package (Macdonald and Du, 2012). Our modal decompositions were considered as valid according to the test from the “mixdist” package, which is based on the χ^2^ approximation to the likelihood ratio statistic. This allowed the identification of gamma components and their parameters – mean, sigma, estimated and proportion – each corresponding respectively to the mean CL size, the standard deviation of the CL size and the proportion of a defined cohort.

### Thermal and Environmental niches of each species, sex and life stage

Four temperature descriptors were calculated for each of the 20 population samples where temperature was measured: the mean (Avg.T), the minimum (Min.T), the maximum (Max.T) and the standard deviation (Std.T) (Table S3). Four additional environmental measures – pH, H_2_S, Fe^2+^ and Fe_tot_ concentrations – were used for 14 of these samples (Table S3) for a second more refined niche analysis. Values below the detection limit of the measurement devices were considered as null for the niche analysis. Pearson’s correlations were used to select non-redundant variables.

The niches of males, females, subadults and juveniles of each shrimp species were examined similarly to the study of thermal niches of species encountered in *B. azoricus* mussel beds at the Lucky Strike vent field (Husson *et al*., 2017). The niche of a set of specimens (*i*.*e*. males, females, subadults or juvenile stages) was studied using the outlier mean index (OMI) created by (Dolédec *et al*., 2000), and computed using the “niche” function of R package ade4 (Dray and Dufour, 2007). On a PCA of environmental parameters, sampling points corresponding to the different population samples collected are weighted by the specimen’s number. The center of gravity of each of these weighted points is the average position of a group of individuals – defined here by species and life stage or sex – in the scatterplot defined by the PCA, *i*.*e*., the average environmental conditions in which this group thrives in. Different indexes are given by the OMI analysis including the OMI index, also called “marginality”. A permutation test is used to check the significance of this index, indicating if species marginality is significantly higher than expected by chance. If this test is not significant, this means that the species distribution is independent of the thermal conditions. The OMI analysis also gives a measure of the niche breath: the tolerance index (Tol), which corresponds to the variance around the centroid. Associated to the tolerance (Tol), the residual tolerance (RTol) indexes, defined as the part of the variance that is not explained by the environmental variables used in the PCA, indicates whether the chosen variables are suitable for the niche analysis. These three index (OMI, Tol and RTol) constitute the total inertia of the niche, and each can be expressed as a percentage of this inertia. At the same time, OMI analysis also identifies the thermal variables that best differentiate the thermal niches of the studied group of individuals.

## Results

### Population structures of *Rimicaris* shrimps at TAG & Snake Pit in February-March 2018

At each vent field, overall size-distributions of the two *Rimicaris* species were drastically different from each other with significantly distinct size-frequency distributions both at TAG (Kolmogorov-Smirnov 2-samples test, D = 0.949, p < 0.001) and at Snake Pit vent fields (Kolmogorov-Smirnov 2-samples test, D = 0.545 p < 0.001). Unlike *R. exoculata, R. chacei* size-distribution was indeed skewed towards the smaller sizes, corresponding to the juveniles and subadult life stages (Figure 1). This results in a significantly higher proportion of immature individuals for *R. chacei* compared to *R. exoculata*, either at TAG (*R. chacei*: 91.9%, *R. exoculata*: 23.8%; χ^2^ = 1036.4, p < 0.001; Table 1) or Snake Pit (*R. chacei*: 52.1%, *R. exoculata*: 32.6%; χ^2^ = 45.74, p < 0.001; Table 1). Interestingly, immature individuals represented a significantly higher proportion of the whole *R. chacei* population at TAG compared to Snake Pit (χ^2^ = 240.3, p < 0.001), whereas immature individuals were significantly more represented in Snake Pit populations of *R. exoculata* compared to TAG (χ^2^ = 34.2, p < 0.001). Nonetheless, a similar proportion of *R. chacei* stage A juveniles were observed in the populations of both vent fields (TAG: 41.2%, Snake Pit: 38.1%; χ^2^ = 0.79, p = 0.373). *R. exoculata* stage A juveniles however, were still in lower proportions at the TAG vent field (TAG: 3.7%, Snake Pit: 13.9%; χ^2^ = 108.03, p < 0.001).

**Table 1.**
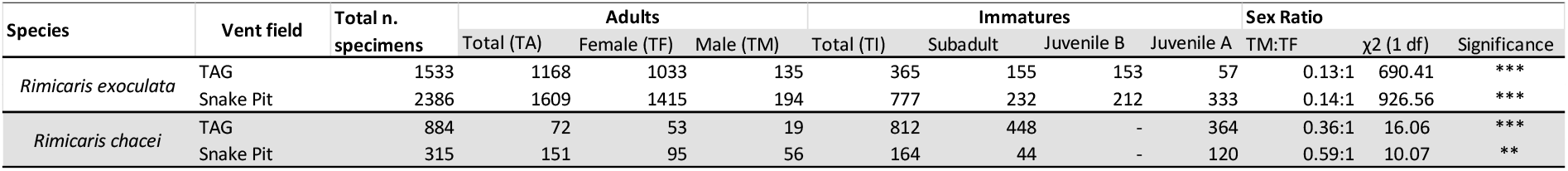
Sample and population data of *Rimicaris exoculata* and *Rimicaris chacei* from TAG and Snake Pit.

**Figure 1.**
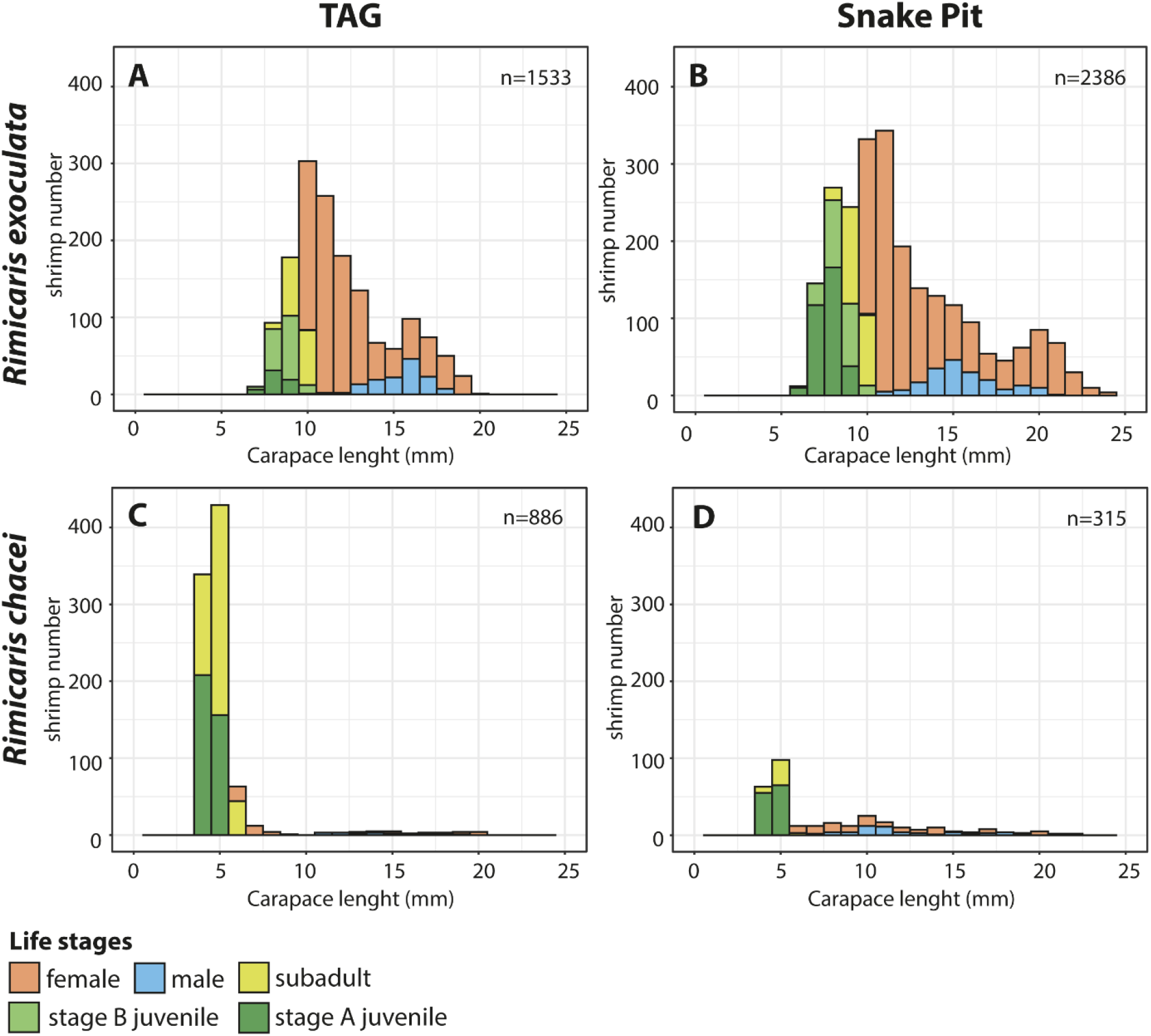
Size-frequency distribution of *Rimicaris exoculata* and *R. chacei* from two vents fields, denoting life stages and sexes. **A**. *R. exoculata* from TAG. **B**. *R. exoculata* from Snake Pit. **C**. *R. chacei* from TAG. **D**. *R. chacei* from Snake Pit.

Overall, larger sizes were observed for *R. exoculata* compared to *R. chacei* at any life stage, either for adults, subadults or juvenile stages (Table S4; Mann-Whitney test, p < 0.001, all cases). Largest recorded adults of the two species were nonetheless of similar size at TAG (20.1 mm and 20.5 mm CL respectively for *R. exoculata* and *R. chacei*) but were bigger for *R. exoculata* at Snake Pit (24.4 mm and 21.1 mm CL respectively for *R. exoculata* and *R. chacei*). Males of *R. exoculata* exhibited significantly larger average sizes than females in both vent fields (TAG: W = 24460, p < 0.001; Snake Pit: W = 80537, p < 0.001). However, the largest recorded *R. exoculata* adults were females both at TAG and at Snake Pit (20.1 mm and 24.4 mm CL respectively). In *R. chacei*, the average size of males and females was not significantly different at Snake Pit (W= 2180, p = 0.065) and males were slightly larger than females in average at TAG (W = 249.5, p = 0.001). Like for *R. exoculata*, largest recorded *R. chacei* adults were females at both vent fields (20.5 mm and 21.8 mm CL respectively).

At each vent field, MIX modal decomposition detected four cohorts in the populations of *R. exoculata* and three cohorts in the populations of *R. chacei* (Figure S2). In *R. exoculata*, the second and third cohorts had the highest proportions of individuals, gathering roughly two third of the populations, both at TAG and Snake Pit (Table 2). These 2 cohorts mostly gathered adults with some immature stages below the OSD in the second cohort. The first cohort only comprised immature individuals and represented at most 1/5 of the total population. The fourth cohort comprised the larger adults, and represented at most 1/5 of the total population. In *R. chacei* however, the first cohort, that gathered mostly immature individuals, comprised half of the population at Snake Pit and up to 92.5% of the population at TAG. The second cohort of *R. chacei* comprised small adults mostly, with proportions varying between 3 to nearly 40% of the whole population. The third cohort comprised large adults representing only 4 to 12% of the total population (Table 2).

**Table 2.** Cohort analysis with proportions (%), average (µ) and standard deviations (σ) of the modal components estimated from the size–frequency distributions of *Rimicaris exoculata* and *Rimicaris chacei* collected from TAG and Snake Pit.

| Species | Vent field | Cohort 1 |  |  | Cohort 2 |  |  | Cohort 3 |  |  | Cohort 4 |  |  |
| --- | --- | --- | --- | --- | --- | --- | --- | --- | --- | --- | --- | --- | --- |
| | | % | $\mu$ | $\sigma$ | % | $\mu$ | $\sigma$ | % | $\mu$ | $\sigma$ | % | $\mu$ | $\sigma$ |
| *Rimicaris exoculata* | TAG | 14.5 | 8.6 | 0.6 | 25.2 | 10.3 | 0.6 | 39.6 | 11.9 | 1.3 | 20.7 | 16.5 | 1.3 |
|  | Snake Pit | 21.7 | 7.9 | 0.7 | 33.5 | 10.5 | 0.9 | 33.6 | 13.7 | 2.4 | 11.2 | 20.4 | 1.4 |
| *Rimicaris chacei* | TAG | 92.5 | 4.7 | 0.4 | 3.3 | 6.8 | 0.3 | 4.2 | 15.3 | 3.2 | - | - | - |
|  | Snake Pit | 48.9 | 4.6 | 0.3 | 39.4 | 9.4 | 2.5 | 11.7 | 17.4 | 2.4 | - | - | - |

Populations of the two *Rimicaris* species differed also slightly by their sex ratio. As in 2014 (Hernández-Ávila *et al*., 2021), sex ratios of *R. exoculata* collected in 2018 from TAG and Snake Pit were strongly biased towards females at both vent fields (TAG: χ^2^ = 690.41, p < 0.001; Snake Pit: χ^2^ = 926.56, p < 0.001; Table 1). Similarly, sex ratios of the *R. chacei* collected at these vent fields deviated significantly from 1:1 (TAG: χ^2^ = 16.06, p < 0.05.; Snake Pit: χ^2^ = 10.07, p < 0.001; Table 1). However, this deviation from a 1:1 sex ratio was less important in *R. chacei* populations compared to *R. exoculata* ones, with a much larger proportion of males among adult individuals at TAG (*R. chacei*: 26.4%, *R. exoculata*: 11.6%; χ^2^ = 12.38, p < 0.001) and at Snake Pit (*R. chacei*: 37.1%, *R. exoculata*: 12.1%; χ^2^ = 68.92, p < 0.001). For each species, sex ratios did not vary significantly between populations of the two vent fields (*R. chacei*: χ^2^ = 0.12, p = 0.732; *R. exoculata*: χ^2^ = 2.04, p = 0.153).

### Spatial variation in the population structure of *Rimicaris* shrimps from TAG & Snake Pit

Our 30 samples were collected into six visually distinct types of assemblages (Figure 2) according to the dominant *Rimicaris* life stage/species/sex and the proximity of the sampled assemblage to the hydrothermal activity. More specifically, these assemblages were classified as close to active emissions for those adjacent to black smokers, close to diffuse emissions for those in a close proximity to transparent shimmering flows, or in inactive areas when no vent fluid could be detected visually in their surroundings. A number of six clusters was also determined statistically to be the most optimal number of clusters for our population dataset using the gap statistic method (Figure 3).

**Figure 2.**
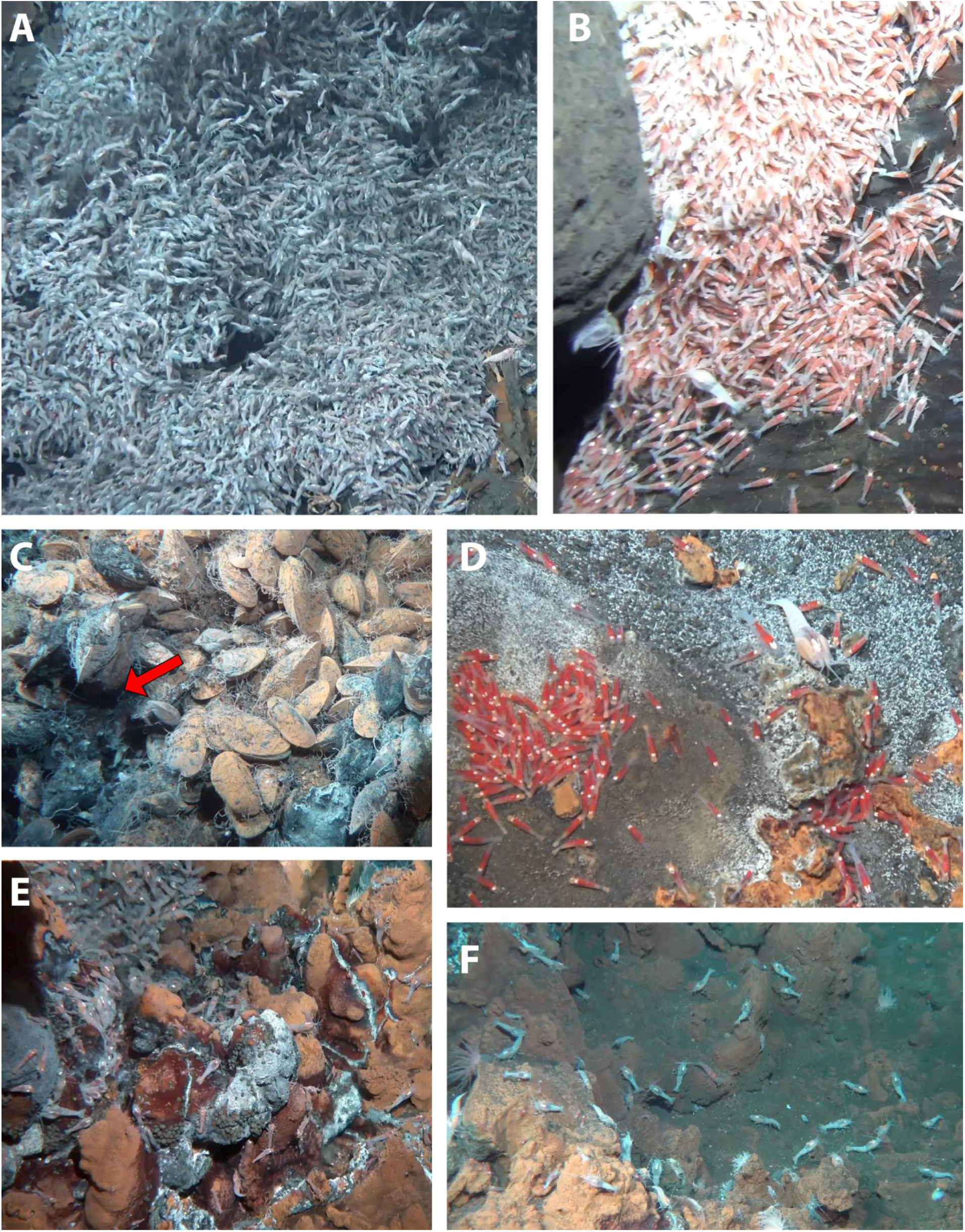
Overview of the different *Rimicaris* assemblage types from TAG and Snake Pit **A**. Dense aggregate of *R. exoculata* adults observed near vent fluid exits. **B**. Nursery of *R. exoculata* adjacent to the dense adult aggregate on the flank of chimneys at Snake Pit. **C**. Hidden aggregate of *R. chacei* observed behind a mussel bed at Snake Pit. **D**. *R. chacei* nursery associated with low temperature diffusions. **E**. Low density alvinocaridid assemblage at the periphery of a dense aggregate of *R. exoculata* at TAG. **F**. Scattered *Rimicaris* individuals, in areas away from any visible vent fluid emissions.

**Figure 3.**
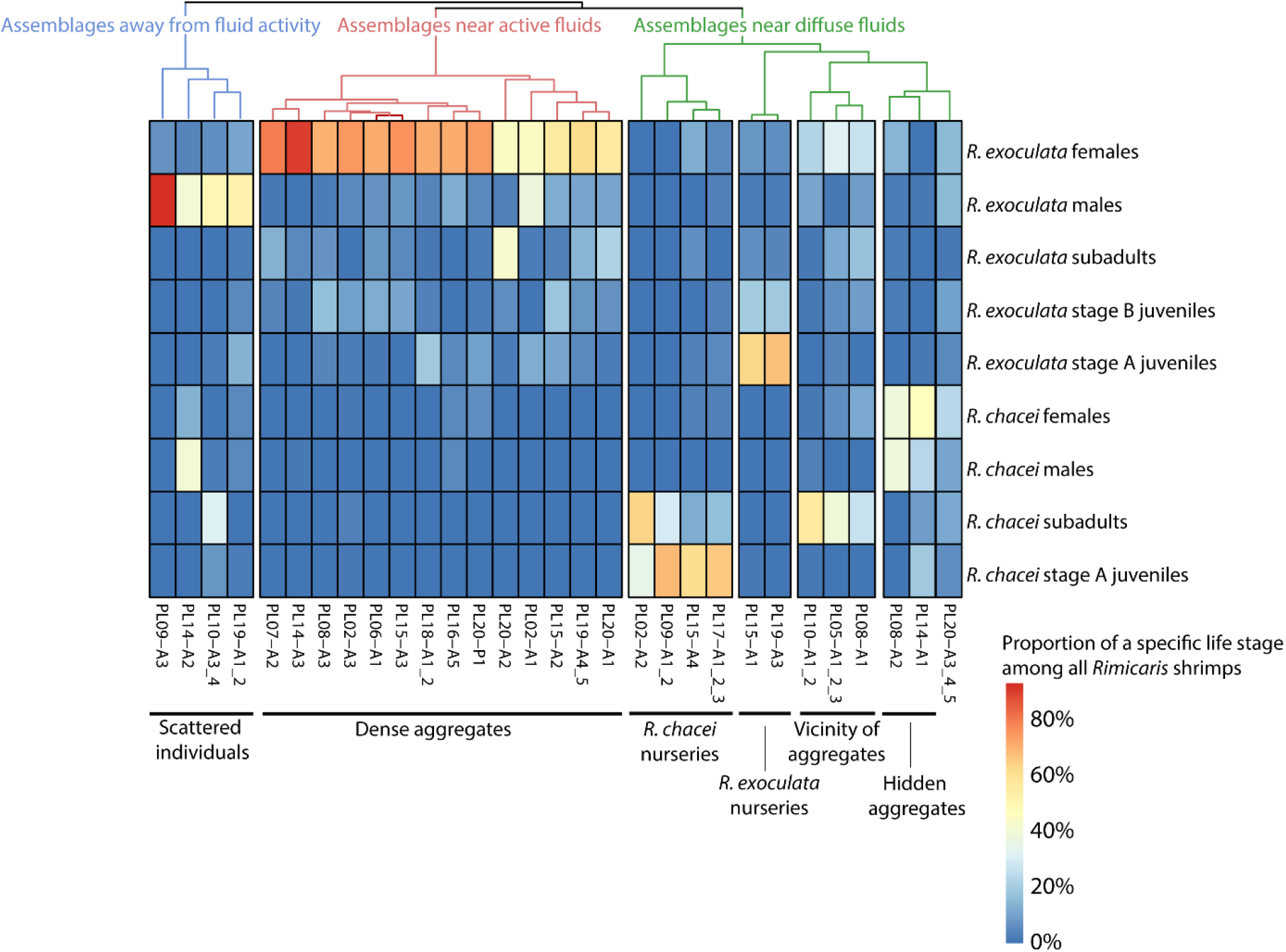
Heat map displaying hierarchical clustering of the different samples collected in this study, according to their composition (in terms of species, sexes and life stages).

The most visually dominating assemblage type was the dense aggregates observed close to the active vent fluid emissions (Figures 2A and S3). These assemblages comprised almost exclusively *R. exoculata*, with *R. chacei* sometimes also present but in very low proportions (*%R. chacei:* 0.9% – 10.4% of all *Rimicaris* shrimps; χ^2^ = 93.5 – 714.1, p < 0.001). Most of the dense aggregates were dominated by adults of *R. exoculata*, with a larger proportion of adults in all samples ((*%Adults:* 67.4% – 94.2% of all *R. exoculata* shrimps; χ^2^ = 47.1 – 501.1, p < 0.001), except in one from Snake Pit (The Nail edifice) that exhibited similar proportions of adults and immature individuals (χ^2^ = 0.46, p = 0. 49). Immatures from this population sample were in majority at an advanced juvenile stage (*i*.*e*., subadults or stage B juvenile), early juveniles only representing 2.6% of all the *R. exoculata* individuals. Large variations were observed in the proportions of immature stages between the different aggregates from 5.8% to 32.6% of all the *R. exoculata* individuals. For nearly all the dense aggregates, sex ratio of *R. exoculata* strongly deviated from 1:1 (χ^2^ = 31.2 – 311, p < 0.001), with a large bias toward females. One exception though, was found in a dense aggregate from TAG exhibiting an equilibrated sex ratio for *R. exoculata* (χ^2^ = 0.69, p > 0.05).

At the periphery of the dense aggregates, in parts less impacted by the turbulent mixing between hydrothermal fluid and seawater, or where fluid exits were dampened, a second type of assemblage was observed at Snake Pit (The Beehive), also largely dominated by *R. exoculata* (Figures 2B and S4A). Unlike dense aggregates, these two population samples exhibited large proportions of immatures (χ^2^ = 202.2, p < 0.001; 231.2, p < 0.001) representing 90.4% and 91.3% of all the *R. exoculata* individuals collected there. These were called nurseries of *R. exoculata*. Most of these immature individuals were at an early juvenile stage, with stage A juvenile representing 70.4% and 74.5% of all the immature individuals. In addition, sex ratio of the few *R. exoculata* adults from these assemblages did not significantly deviate from 1:1 (χ^2^ = 3.2, p = 0.07) or only slightly toward females (χ^2^ = 5.4, p = 0.02). As for the dense aggregates, *R. chacei* shrimps were nearly absent from *R. exoculata* nurseries with only two *R. chacei* juveniles collected in total.

Two other types of shrimp assemblages observed close to diffuse vent fluid emissions were dominated by *R. chacei* (χ^2^ = 16.5 – 888.1, p < 0.001) representing 77.4% to 99.5% of all the *Rimicaris* shrimps collected. These were generally not directly adjacent to the dense aggregates, but clearly separated even if the distance was not necessarily important. One of these two types of assemblages was collected in a hole between rocks at TAG or between *Bathymodiolus puteoserpentis* mussels at Snake Pit (The Moose edifice). Although individuals were rather densely packed, they were hard to see due to the relatively small size of the assemblages and their “hidden” localization. These assemblages were called “hidden aggregates” of *R. chacei* (Figure 2C and S4B). These hidden aggregates clearly exhibited larger proportions of *R. chacei* adults than immatures (χ^2^ = 35.6 – 60.1, p < 0.001), adults representing 71.7% to all the *R. chacei* individuals collected there. Sex ratio of *R. chacei* from these assemblages was strictly equilibrated for the sample collected at TAG (χ^2^ = 0, p = 0.99) but slightly deviated toward females for the one collected at Snake Pit (χ^2^ = 7.45, p = 0.006).

The second type of assemblages dominated by *R. chacei*, was observed close to diffuse emissions exiting from cracks at the bottom of vent chimneys or in flat areas of the vent fields (Figures 2D and S4C). Contrary to the hidden aggregates, significantly larger proportions of immature individuals were observed in these nurseries of *R. chacei* (χ^2^ = 36.8 – 868.5, p < 0.001), representing at least 96.7% of all the individuals of the species. In general, most of these immature individuals were at an early juvenile stage, with stage A juvenile representing 69.9% to 82.6% of all the immature individuals. Though, one *R. chacei* nursery from TAG exhibited a larger proportion of *R. chacei* subadults than stage A juveniles.

A last assemblage type found in areas of moderate fluid influence was observed at TAG in the vicinity of dense aggregates (Figures 2E and S4D). These assemblages were characterized by similar proportions of the two *Rimicaris* species (χ^2^ = 2.9, p > 0.05) or only slightly larger proportions of one of the two species (χ^2^ = 7.5 – 7.9, p < 0.05). Larger proportions of *R. exoculata* adults (χ^2^ = 24 – 30.1, p < 0.001) and larger proportions of *R. chacei* immatures (χ^2^ = 56.4 – 208.9, p < 0.001) were usually observed in these assemblages. However, in one of these assemblage, similar proportions of adults and immature individuals were found for both species (χ^2^ = 0.4 – 2.5, p > 0.05). In all of them, most of the immature shrimps were at an advanced juvenile stage for both species, stage A juveniles representing at most 7.3% and 2.4% of all the immature individuals respectively for *R. exoculata* and for *R. chacei*. In general, sex ratios of *R. exoculata* from these assemblages did not significantly deviate from 1:1 (χ^2^ = 0.1 – 3, p > 0.05), except in one with a large bias toward females (χ^2^ = 117.4, p < 0.001).

At last, *Rimicaris* shrimps were also collected at the periphery of the vent chimneys, on inactive sulfides away from the vent fluid activity (Figures 2F and S4E). These assemblages of scattered individuals exhibited either similar proportions of the two *Rimicaris* species (χ^2^ = 0.1 – 2, p > 0.05) or larger proportions of *R. exoculata* individuals (χ^2^ = 26.4 p < 0.001). As in dense aggregates, larger proportions of *R. exoculata* adults over immature individuals were observed in all of these assemblages (χ^2^ = 16.2 – 44.6, p < 0.001). Sex ratio of *R. exoculata* from these assemblages deviated from 1:1 (χ^2^ = 6.4 – 16.3, p < 0.001) with a large bias toward males. For *R. chacei* both larger proportion of adults (χ^2^ = 20.2, p < 0.001) and larger proportions of immature individuals (χ^2^ = 28.9, p < 0.001) were reported in these two different population samples. Unlike *R. exoculata*, sex ratio of *R. chacei* from these assemblages did not significantly deviate from 1:1 (χ^2^ = 3, p = 0.11).

One population sample, collected in a flat area with moderate fluid emission away from an active venting chimney at Snake Pit (The Nail edifice), did not exhibited a clear structure corresponding to one of the six assemblages defined above. It rather clustered with the hidden aggregates of *R. chacei* adults (Figure 3). However, unlike samples from hidden aggregates, this population showed similar proportions of both *Rimicaris* species (χ^2^ = 0.2, p > 0.05). Moreover, proportion of adults and immature individuals were similar (χ^2^ = 0.1 – 1.4, p > 0.05) and sex ratio did not significantly deviate from 1:1 (χ^2^ = 0.1 – 0.11, p > 0.05) both for *R. exoculata* and *R. chacei*. This assemblage also differed from the others by being dominated by *Mirocaris fortunata* (58.9% of all the shrimps sampled), another co-occurring alvinocaridid at these sites that was sporadically observed in very low proportions in the other samples.

### Thermal & environmental niches of *Rimicaris* shrimps and their life stages from TAG & Snake Pit

Average temperatures around *Rimicaris* assemblages were comprised between 3.4°C and 22.5°C in average, with temperature reaching up to 38.1°C and minimum records down to 3.1°C. Standard deviations that can be considered as the thermal variability around the assemblages varied from 0.3°C to 6.9°C. Regarding the vent fluid chemistry, H_2_S concentrations around *Rimicaris* assemblages were comprised between 0 µM and 8.5 µM in average with pH variations between 6.86 and 7.9 as well. It is worth mentioning that total sulfides concentrations were measured onboard on recovered samples and may be underestimated compared to the actual *in situ* conditions. *In situ* iron concentrations around *Rimicaris* assemblages were highly variables with Fe^2+^ concentrations comprised between 0 µM and 239 µM and Fe_tot_ concentrations between 0 µM and 316 µM. Concerning the PCA analysis, minimal temperatures were strongly correlated with average temperature (r^2^ = 0.97, p < 0.05), only average temperature (Avg.T), temperature standard deviation (Std.T) and maximum temperature (Max.T) were used in the two PCA. No clear significant correlation was observed between any of the thermal descriptors and the concentrations of the different reduced chemical elements (Pearson correlations, p > 0.01 in all cases).

For the thermal niches (Figure 3A and 3B; 20 sampling points), most of the variance was explained by the two first axes of the PCA representing respectively 93.6% and 5.3% of the inertia. All three-temperature metrics were very well represented on the PCA (∑cos^2^ = 0.99, 0.99 and 0.98 for Avg.T, Std.T and Max.T respectively) and were strongly correlated with the first axis (cor = 0.96, 0.95 and 0.99 for Avg.T, Std.T and Max.T respectively). Avg.T, Std.T and Max.T almost contributed equally to the construction of the first axis (contribution = 32.7%, 32.4% and 34.9% respectively). Similarly, Avg.T and Std.T also contributed equally to the construction of the second axis (contribution = 46.4% and 53.4% respectively).

In the case of the environmental niches (Figure 3C and 3D; 14 sampling points), a large part of the variance was explained by the two first axes of the PCA representing respectively 64.7% and 22.4% of the inertia. All the environmental parameters were relatively well represented on the two first axes of the PCA with ∑cos^2^ > 0.85 for all parameters except for Std.T and pH (∑cos^2^ = 0.81 and 0.70 respectively). Thermal descriptors as well as pH correlated more with the first axis (cor = −0.97, −0.88, −0.95 and 0.83 for Avg.T, Std.T, Max.T and pH respectively) whereas H_2_S measurements were more correlated with the second axis (cor = −0.83). Iron concentrations were equally correlated with the first (cor = −0.75 and −0.67 for Fe^2+^ and Fe_tot_ respectively) and the second axis of the PCA (cor = 0.62 and 0.64 for Fe^2+^ and Fe_tot_ respectively). The thermal descriptors contributed the most to the construction of the first axis (contribution = 21%, 17% and 20.1% respectively for Avg.T, Std.T and Max.T), followed by pH and Fe^2+^ and Fe_tot_ (contribution = 15.4%, 12.5% and 10% respectively). On the other hand, H_2_S contributed in large part to the construction of the second axis followed by iron concentrations (contribution = 24.3% and 25.8% respectively for Fe^2+^ and Fe_tot_).

As revealed by the OMI analysis, stage A juveniles of *R. exoculata* and *R. chacei* occupied distinct thermal niches compared to their older counterparts (Figure 4A). The thermal niches of these early life stages differed both in terms of niche breadth and overall position (Figure 4B). Indeed, stage A juveniles of *R. exoculata* exhibited a much lower tolerance (Tol) than the other adults or immature stages of this species (Figure 4B). Similarly, stage A juveniles of *R. chacei* also displayed a lower tolerance (Tol) than the other *R. chacei* life stages, as well as a mean niche position that significantly deviated from the average sampled thermal conditions (Permutation test, p < 0.05). Additionally, permutation tests also showed a significant marginality for *R. exoculata* females and subadult stages (p < 0.05; Table 3) despite a rather similar tolerance (Tol) compared to *R. exoculata* males and stage B juveniles (Table 3). Conversely, niche position of *R. chacei* adults and subadults did not significantly departed from the average sampled thermal conditions. Overall, every *R. exoculata* life stages presented a larger niche breadth than any of the *R. chacei* life stages (Figure 4A). The same trends were also observed in the more detailed environmental niches with distinct niches for the earliest juvenile stages of both *R. exoculata* and *R. chacei*, compared to all the other life stages (Figure 4C and 4D) and a lower niche breadth for *R. chacei* compared to any *R. exoculata* life stages (Figure 4C and 4D), although significant marginality was only found for *R. exoculata* females (p < 0.05; Table 3). For the thermal niches, residual tolerance (RTol) ranged from 0.3% to 10.2% of total inertia; thus, the variables used to describe these niches explained 89.8% to 99.7% of the niche distribution of the different *Rimicaris* life stages (Table 3). On the other hand, residual tolerance (RTol) for the environmental niches ranged from 26.6% to 49% of total inertia; thus, the variables used to describe these niches only explained 51% to 74.4% of the niche distribution of the different *Rimicaris* life stages (Table 3).

**Table 3.**
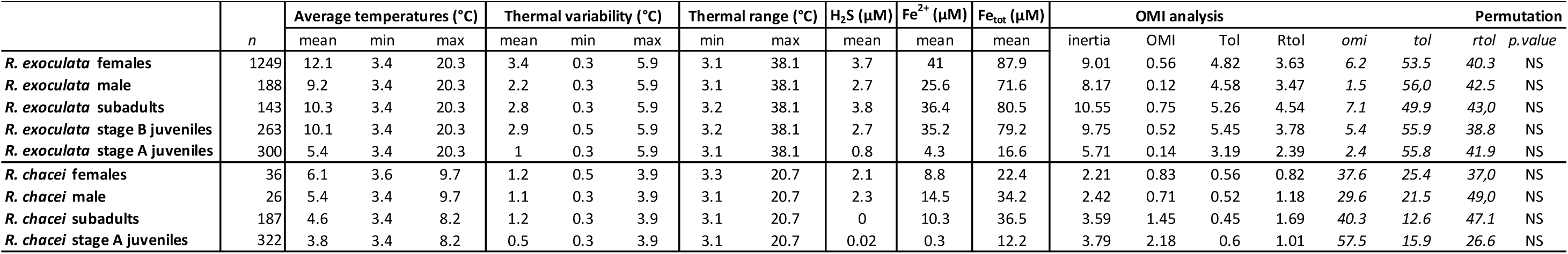
Thermal range and parameters of the OMI analysis for the different *Rimicaris* groups.

**Figure 4.**
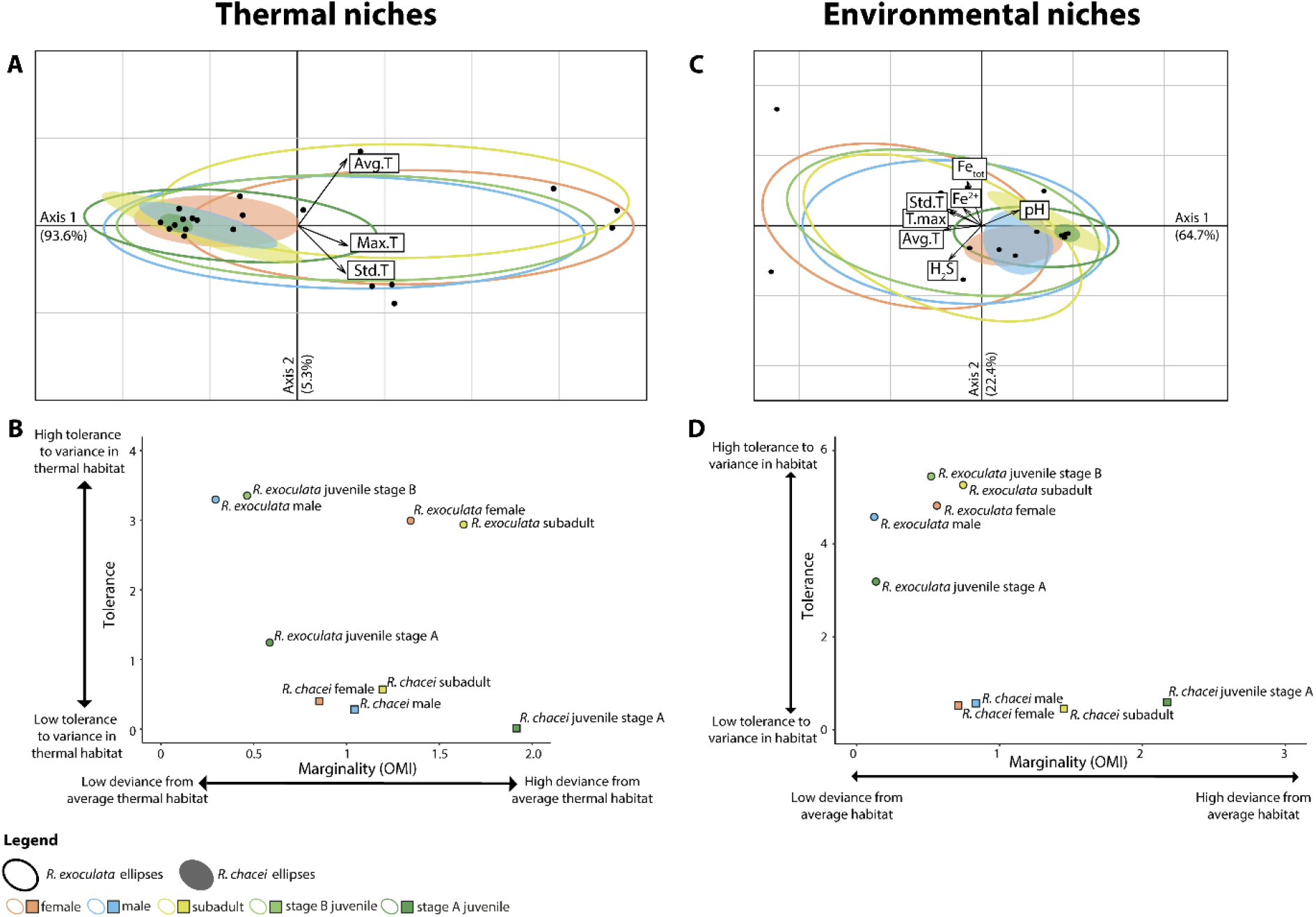
Niches of *Rimicaris* shrimps according to their species, sex and life stages. **A**. Two-dimensional plot of the OMI analysis. Each ellipse represents the thermal niche of a *Rimicaris* group (defined by its species, life stage, and sex in adults). **B**. *Rimicaris* groups total marginality (OMI index) versus total tolerance of their thermal niches. **C**. Two-dimensional plot of the OMI analysis. Each ellipse represents the environmental niche of a *Rimicaris* group (defined by its species, life stage, and sex in adults). **D**. *Rimicaris* groups total marginality (OMI index) versus total tolerance of their environmental niches.

## Discussion

### Small scale distribution of *Rimicaris* life stages along the environmental gradient

Our temperature measurements indicate that thermal conditions around the different *R. exoculata* assemblages of TAG and Snake Pit are comprised between 3.4°C and 22.5°C with maximal peaks of temperatures up to 38.1°C (Table 3). These results slightly extend the known thermal range occupied by these shrimps compared to previous *in situ* temperature measurements around *R. exoculata* aggregates, which reported average thermal conditions between 4.6°C and 16°C and up to 25°C at maximum (Desbruyères *et al*., 2001; Geret *et al*., 2002; Zbinden *et al*., 2004; Schmidt, Le Bris and Gaill, 2008). Although *R. exoculata* reaches 100% of mortality when exposed during 1h at 33°C only in closed pressurized chambers, these shrimps tolerate better shorter exposures of 10 min with only 35% of mortality at 39°C (Ravaux *et al*., 2019). With a defined critical thermal maximum (CTmax) of 38°C for this species, it thus appears that *R. exoculata* occupies *in situ* most of its fundamental thermal niche. We also noticed important movements of shrimps in aggregates where average temperature exceed 20°C – PL06A1, PL15A3 & PL20A1 – whereas the others tend to remain steady confirming the *in vivo* behavioral observations of Ravaux et al. (Ravaux *et al*., 2019). On the other hand, we observed that *R. chacei* inhabits a narrower thermal niche, comprised between 3.4°C and 14.5°C – up to 20.7 °C at maximum – whatever the life stage considered (Table 3). In the absence of experimental work on *R. chacei* thermotolerance, it is unclear if their more limited distribution within the hydrothermal vent gradient stems from a physiological limitation or from biological interactions such as competition with its congeneric species. Part of the answer can be brought by a comparison with *M. fortunata*, another alvinocaridid shrimp from the MAR, with a thermal tolerance similar to *R. exoculata* (Shillito *et al*., 2006) but occupying a more restricted thermal habitat (Desbruyères *et al*., 2001; Methou, 2019). The narrower distribution of *R. chacei* could also simply be related to a lower thermal preference, as seen in many vent species that preferentially selects habitats with cooler temperatures than their upper thermal limit (Bates *et al*., 2010). This all suggests that temperature *per se* is not a sufficient limiting factor to explain differences in alvinocaridid shrimp distributions.

Similarly, our dataset also points out that *R. exoculata* inhabits in a wider range of reduced compound concentrations (traced through *in situ* total dissolvable iron) than *R. chacei* for a given life stage, although this trend was much less marked between the earliest juvenile stages of the two species. Our *in situ* chemistry measurements around biological communities indicate significant deviation from the conservative dilution model as it has already been reported in previous studies (Le Bris *et al*., 2006, 2019; Nakamura and Takai, 2014). This deviation from a conservative dilution of the endmember originates from the highly reactive nature of the sulfides and iron species and the consumption or production of these reduced compounds by microbial activities.

At both fields, an important zonation between adults and juvenile stages was also found for *R. exoculata* and for *R. chacei*. This spatial segregation along the thermal gradient is reflected in the distinct thermal niches observed between the earliest juveniles and the other life stages for the two *Rimicaris* species. Similar differences in distribution of bathymodiolin mussel life stages have been hypothesized to be related to intraspecific competition for space between adults and juveniles (Comtet and Desbruyères, 1998; Husson *et al*., 2017). This was further supported by evidence of recruitment inhibition by high density assemblages of adult mussels from the East Pacific Rise (Lenihan *et al*., 2008). Still, an influence of intraspecific competitive interactions among *Rimicaris* shrimps is tempered by their high motility as well as the non-negligible presence of early juveniles within several assemblages of high densities dominated by adult stages. A progressive acquisition of a high thermal tolerance of these recruiting stages, as seen in post-larvae stages of the intertidal shrimp *Palaemon elegans* (Ravaux *et al*., 2016), could also limit their upper thermal distribution. Additionally, distribution of *R. exoculata* and *R. chacei* juveniles further differ by their habitat topography, on steep chimney walls for *R. exoculata* nurseries and within flat areas or smooth slopes for *R. chacei* nurseries. As topography strongly influences the overall current hydrodynamics, with low topography areas generally associated with low mixing rates (Girard *et al*., 2020), these structural differences in habitats might reflect the relative dependency of each species to the vent fluid mixing gradient essential to fuel their ongoing developing symbiosis (Methou *et al*., 2020). Although our measurements were limited, this variability of the fluid chemistry between the nursery habitats of the two species could be observed in our dataset with lower H_2_S and Fe concentrations in *R. chacei* nurseries compared to *R. exoculata* ones.

We also confirmed the segregation pattern between males and females of *R. exoculata* observed at TAG in January-February 2014 (Hernández-Ávila *et al*., 2021) for both of the studied vent fields. Such pattern was not observed in *R. chacei*, which also exhibited a more equilibrated sex ratio overall, more similar to what has been observed for its sister species *R. hybisae* (Nye, Copley and Tyler, 2013).

Despite an apparent relative stability with the maintained presence of an assemblage type in the same area over the years, the spatial zonation of the different *Rimicaris* life stages also seems dynamic with a succession of assemblages in other areas on short temporal scales. During one of our dives, we witnessed at TAG, the entire disappearance of a dense aggregation assemblage of *R. exoculata* adults replaced by an aggregate of *R. chacei* hidden between rocks in just about ten days (Figure S5). Unfortunately, we are currently unable to confirm if this succession of assemblages is related to a modification of the environmental conditions, as we could not obtain temperature measurements in the first place, prior to the replacement of the dense aggregate, due to technical difficulties during the dive.

In addition to the assemblages dominated by a particular life stage of one of the two *Rimicaris* species, different types of mixed assemblages were also observed in the areas a little further away from the fluids. Interestingly, these assemblages with a mixed proportion of the two species were also the assemblages with the lowest densities of *Rimicaris* individuals, visually countable in contrast to the other more dense assemblage types (Figure 2.). These more peripheral assemblages with less marked structuration could correspond to assemblages with much lower densities due to suboptimal habitat conditions for *Rimicaris* spp. Lower supply of reduced compounds to feed their cephalothoracic symbionts, or an absence of a symbiont pool to renew their symbiotic communities along their molt cycle would limit the establishment of these species in these areas. A finer environmental characterization of these particular areas would be required to better understand the subtle ecological differences that could exist between these habitats and the denser shrimp assemblage types. In any cases, these density dependent trends suggest that competitive processes might occur between *R. chacei* and *R. exoculata* life stages.

### Population dynamics of *Rimicaris* shrimps from the MAR are driven by biotic interactions

As most species from hydrothermal vents, *R. exoculata* and *R. chacei* exhibited multimodal size-frequency distribution suggesting a discontinuous recruitment pattern in both species, thus confirming the previous observations made for *R. exoculata* in 2014 (Hernández-Ávila *et al*., 2021). Still, population structures of the two congeneric *Rimicaris* species from the Mid Atlantic Ridge differed widely with a large post recruitment collapse of *R. chacei* populations at both vent fields. This collapse was even more pronounced for the TAG population where immature individuals represented more than 90% of all the individuals collected. This implies either a high mortality rate of *R. chacei* early post-settlement stages in comparison to *R. exoculata* and/or a large variability in the magnitude of the different larval settlement events. Although temporal tracking of these populations could not be performed, important aggregations of *R. chacei* juveniles in nurseries were already reported between January and February 2014 (Hernández-Ávila *et al*., 2021) as well as between March and April 2017 (Methou *et al*., 2020). It is then unlikely that the observed variations between *R. chacei* life stages abundance would be related to variations in larval supply and/or settlement success between years.

Therefore, we hypothesize that the post recruitment collapse observed in *R. chacei* populations results more probably from a higher juvenile mortality that could be explained by different factors, not necessarily mutually exclusive from each other (Figure 5). Among them, predation could have a major impact on *R. chacei* demography, as it has been recognized as one of the most important cause of juvenile mortalities in shallow water ecosystems (Gosselin and Qian, 1997). The more peripheral and exposed distribution of *R. chacei* juveniles within the vent fields could expose them to a higher predation pressure by populations of *Maractis rimicarivora* anemones found in these areas and particularly abundant at TAG (Copley, Jorgensen and Sohn, 2007). Indeed, we could witness the catch of one juvenile from the *R. chacei* nursery area by these anemones during a dive at TAG in 2014 (Supplementary video material 1). Predation by these carnivorous anemones on *R. exoculata* adults has already been observed (Van Dover *et al*., 1997) and was proposed to affect male individuals from the peripheral areas (Hernández-Ávila *et al*., 2021) which could explain, at least partially, the large sex ratio bias toward females of this species. Additionally, a larger vulnerability to predation by macrourid fishes could also affect preferentially *R. chacei* juveniles. Despite a limited observation time on these areas, we reported twice predation events by these fishes within or close to *R. chacei* nurseries assemblages during our samplings at TAG, both in 2014 and in 2018 (Supplementary video material S1). This higher predatory pressure at TAG would also be in agreement with the greater collapse observed in the populations of *R. chacei* at this site compared to those from Snake Pit (Figure 1.). Contrastingly, the distribution of *R. exoculata* early juveniles, sheltered on the flank of large chimney structures or within the dense aggregates of *R. exoculata* adults probably protect them from this high predatory pressure.

**Figure 5.**
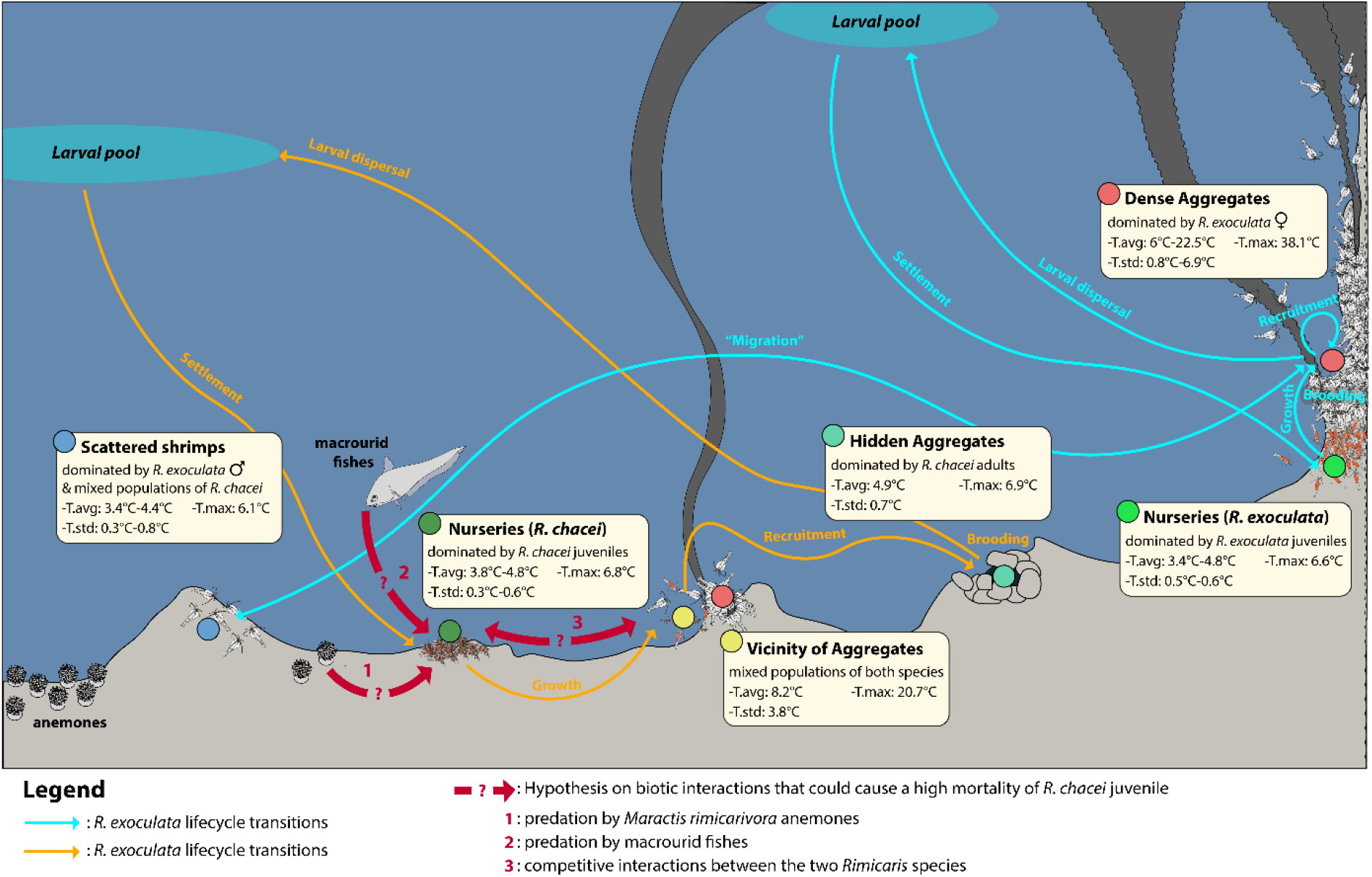
Schematic view of the different Rimicaris assemblage distribution at hydrothermal vents from the Mid Atlantic Ridge with current hypotheses on the factors affecting their population dynamics.

In addition to their distinct spatial distribution, the juvenile stages of both *Rimicaris* shrimps differ in their average settlement size, twice as large for *R. exoculata* compared to *R. chacei* (Methou *et al*., 2020). Numerous studies on marine fishes and shrimps have reported body size as a key factor in the survival of early juveniles with a lower overall mortality for larger individuals (Meekan *et al*., 2006; Gagliano and McCormick, 2007; Mace and Rozas, 2018). Thus, survival of *Rimicaris* juveniles could fit into the “bigger is better” hypothesis, whether larger body size would provide them a lower vulnerability against predation, a better physiological state and/or a higher competitive ability.

As an alternative to a high juvenile mortality, important emigration of *R. chacei* juveniles – secondary migration – in a different location following their initial settlement at the vent fields could also lead to similar patterns in their population structure. A long-term monitoring with video imagery of the areas of *Rimicaris* nurseries or individual tracking early-stage shrimps could help to refine our hypotheses and determine more precisely the relative influence of these different factors on their population dynamics.

Interspecific competition for space between the two species, already suggested by their spatial segregation patterns, could also have a major influence on their demography. It is commonly accepted that abundant species within an assemblage outcompete less common ones. This general statement, however, has been contradicted by various manipulative studies which did not report any effect of being naturally rare or common on the competitive ability of a species (Angel *et al*., 2006; Matias *et al*., 2012). Still, marine lobsters can experience important demographic bottleneck before recruitment, due to the more specific requirements of their early benthic juvenile stages for sheltered areas, which are limited in space (Wahle and Steneck, 1991). In a similar fashion, the low abundance of *R. chacei* adults might result from a lower competitive ability of their early juvenile stages to access areas of active fluid emissions, as suggested by their narrower thermal niche compared to similar stage of *R. exoculata* (Figure 4.).

In hydrothermal vents, competitive processes for space are confounded with competition for food in symbiotrophs whose nutrition depend entirely on their access to the vent fluid reduced elements. Both *R. exoculata* and *R. chacei* are fueled by large communities of filamentous ectosymbionts (Ponsard *et al*., 2013; Apremont *et al*., 2018), which progressively make up during their ontogeny an increasing part of their diet after recruitment (Methou *et al*., 2020). A more limited access to areas of active fluids for *R. chacei* juveniles could potentially lead to a limitation to primarily acquire their episymbionts and develop a sufficient nutritive symbiotic community within their cephalothorax, which could thus cause mass mortality in these populations. In this scenario, the mixotrophic behavior of *R. chacei* would be the result of a past competition with *R. exoculata* leading to a trophic niche differentiation in *R. chacei* adults over evolutionary timescales, allowing the stable coexistence of the two species in the northern MAR vent fields. Additionally, the more limited local diversity at TAG (Desbruyères *et al*., 2001), likely related to the absence of *Bathymodiolus* mussel beds known to house large macrofaunal diversity (Desbruyères *et al*., 2001; Rybakova and Galkin, 2015), enhance even more the limitation in alternative food resources for *R. chacei*, thus possibly explaining the larger post recruitment collapse of their population at this site. These shrimps are not the first case of population demographics driven by competitive interactions for food in hydrothermal vent ecosystems. Populations of *Paralvinella pandorae* from Juan de Fuca Ridge exhibited indeed similar post settlement collapses when found in sympatry with their congeneric species *P. palmiformis*, whose isotopic niches largely overlap (Levesque, Juniper and Marcus, 2003).

This link between nutrition modes and population demographics of these shrimps is also further supported when their population structures are compared with those of *R. hybisae* from the Mid Cayman Spreading Center. The latter is extremely close phylogenetically to *R. chacei*, insofar as the two species cannot be distinguished on the sole basis of the genetic markers used so far (Vereshchaka, Kulagin and Lunina, 2015). In this way, *R. hybisae* populations from the Mid Cayman Rise can almost be seen as a case of *R. chacei* allopatry with no known competitors like *R. exoculata*. However, *R. hybisae* size-frequency structures largely differ from those of *R. chacei* and appear to be more similar to those of *R. exoculata* (Nye, Copley and Tyler, 2013).

To be noticed, unlike *R. exoculata, R. chacei* adults were more difficult to catch with the suction sampler during our attempts to collect them as they were sometimes escaping the tip of the sampler when we approached them. This escape behavior was not observed for the juvenile specimens of the same species inhabiting the nurseries. It is therefore possible that our dataset might slightly underestimate the exact proportion of *R. chacei* adults in the field. Nonetheless, it is unlikely that this potential underestimation of *R. chacei* adult number would overturn the extremely marked population structure observed for this species at both vent fields. Similarly, we believe that the disequilibria in sampling effort between the different assemblage types, with an overrepresentation of the dense aggregates against the other assemblage types, is representative of their higher occurrence within the hydrothermal vent fields. A detailed cartography giving the distribution of these assemblages, with shrimp densities estimations, when possible, over the entire vent fields would help to better assess our dataset representativeness and if our sampling strategy could have affected partially our conclusions on the population dynamics of these two species.

The mechanisms by which *R. chacei* populations maintain a high supply of settlers despite a limited adult number comparatively to *R. exoculata* are still unclear. Larger *R. chacei* populations from yet unknown vent fields in the MAR representing source populations – with TAG and Snake Pit as sink populations – at the metapopulation level or differences in reproductive output between *R. exoculata* and *R. chacei* are possible. Additional work on the reproductive biology of *R. chacei* will be needed to better understand differences in the life history traits of these species. Further experimental studies on the thermal physiology and tolerance to the vent fluid toxic compounds of both *R. chacei* adults and *Rimicaris* juvenile stages would also be required to understand if the distribution patterns observed between these different life stages are related to physiological limitations or other biotic factors.

## Author’s contributions

P.M. collected and sorted the specimens, carried out the measures and identification of specimens, lead data analysis, participated in the conception and design of the study and drafted the manuscript; I.H-A. helped to collect and sort the specimens and critically revised the manuscript; C.C. carried out the collection of environmental measurement in the field, provide her help in the analysis of these data and critically revised the manuscript; M-A.C-B. coordinated the study and helped draft the manuscript; F.P. helped to collect and sort the specimens, conceived and designed the study, coordinated the study and helped draft the manuscript. All authors gave final approval for publication and agreed to be held accountable for the work performed therein.

## Acknowledgements

The authors thank the captains and crews of the R/V *Pourquoi pas*? and the Nautile submersible team for their efficiency, as well as the chief scientist and scientific parties of and BICOSE 2 cruise (http://dx.doi.org/10.17600/18000004). Further thanks go to Leela Morzina for her help in collecting data on shrimp measurements and identification of their life stages and to Emmanuelle Omnes (ifremer, LEP) for her help in sorting animal samples on board. The use of the CHEMINI device was possible thanks to the help of Dr. Agathe Laës-Huon (ifremer, LDCM). Maps of the sampling sites was provided by Anne-Sophie Alix and Florian Besson (Ifremer, CTDI). We also thanks Lyndsay Clavareau (ifremer, DYNECO) for her help to carry out the niche analysis. This work was supported by the Ifremer REMIMA program and the Region Bretagne ARED funding. The authors declare they have no competing interests.

## Literature cited

Angel, A. et al. (2006) ‘Causes of rarity and range restriction of an endangered, endemic limpet, Siphonaria compressa’, Journal of Experimental Marine Biology and Ecology, 330(1), pp. 245–260. doi: 10.1016/j.jembe.2005.12.031.

Apremont, V. et al. (2018) ‘Gill chamber and gut microbial communities of the hydrothermal shrimp Rimicaris chacei Williams and Rona 1986: A possible symbiosis’, PLoS ONE. edited by C.-H. Kuo, 13(11), p. e0206084. doi: 10.1371/journal.pone.0206084.

Bates, A. E. et al. (2010) ‘Deep-sea hydrothermal vent animals seek cool fluids in a highly variable thermal environment’, Nature Communications, 1(May), pp. 1–6. doi: 10.1038/ncomms1014.

Beermann, J. and Franke, H. D. (2012) ‘Differences in resource utilization and behaviour between coexisting Jassa species (Crustacea, Amphipoda)’, Marine Biology, 159(5), pp. 951– 957. doi: 10.1007/s00227-011-1872-7.

Bouchemousse, S., Lévêque, L. and Viard, F. (2016) ‘Do settlement dynamics influence competitive interactions between an alien tunicate and its native congener?’, Ecology and Evolution, 7(1), pp. 200–213. doi: 10.1002/ece3.2655.

Le Bris, N. et al. (2000) ‘A new chemical analyzer for in situ measurement of nitrate and total sulfide over hydrothermal vent biological communities’, Marine Chemistry, 72(1), pp. 1–15. doi: 10.1016/S0304-4203(00)00057-8.

Le Bris, N. et al. (2006) ‘Is temperature a good proxy for sulfide in hydrothermal vent habitats?’, Cahiers de Biologie Marine, 47(4), pp. 465–470.

Le Bris, N. et al. (2019) ‘Hydrothermal Energy Transfer and Organic Carbon Production at the Deep Seafloor’, Frontiers in Marine Science, 5(January), pp. 1–24. doi: 10.3389/fmars.2018.00531.

Cline, J. D. (1969) ‘Spectrophotometric determination fo hydrogen sulfide in natural waters’, Limnology and Oceanography, 14(3), pp. 454–458.

Comtet, T. and Desbruyères, D. (1998) ‘Population structure and recruitment in mytilid bivalves from the Lucky Strike and Menez Gwen hydrothermal vent fields (37°17’ N and 37°50’ N on the Mid-Atlantic Ridge)’, Marine Ecology Progress Series, 163, pp. 165–177. doi: 10.3354/meps163165.

Copley, J. T. P., Jorgensen, P. B. K. and Sohn, R. A. (2007) ‘Assessment of decadal-scale ecological change at a deep Mid-Atlantic hydrothermal vent and reproductive time-series in the shrimp Rimicaris exoculata’, Journal of the Marine Biological Association, 87, pp. 859– 867. doi: 10.1017/S0025315407056512.

Desbruyères, D. et al. (2001) ‘Variations in deep-sea hydrothermal vent communities on the Mid-Atlantic Ridge near the Azores plateau’, Deep-Sea Research Part I: Oceanographic Research Papers, 48(5), pp. 1325–1346. doi: 10.1016/S0967-0637(00)00083-2.

Dolédec, S. et al. (2000) ‘Niche separation in community analysis: A new method’, Ecology, 81(10), pp. 2914–2927. doi: 10.1890/0012-9658(2000)081[2914:NSICAA]2.0.CO;2.

Van Dover, C. L. et al. (1997) ‘Predatory Anemones at TAG’, BRIDGE Newsletter, 12.

Van Dover, C. L. (2019) ‘Inactive Sulfide Ecosystems in the Deep Sea:A Review’, Frontiers in Marine Science, 6(July), pp. 1–40. doi: 10.3389/fmars.2019.00461.

Dray, S. and Dufour, A.-B. (2007) ‘The ade4 Package: Implementing the Duality Diagram for Ecologists’, Journal of Statistical Software, 22(4), pp. 1–19. doi: 10.18637/jss.v022.i04.

Faure, B. et al. (2007) ‘Spatial and temporal dynamics of reproduction and settlement in the Pompeii worm Alvinella pompejana (Polychaeta: Alvinellidae)’, Marine Ecology Progress Series, 348, pp. 197–211. doi: 10.3354/meps07021.

Fornari, D. J. D. J. et al. (2012) ‘The East Pacific Rise between 9°N and 10°N: Twenty-five years of integrated, multidisciplinary oceanic spreading center studies’, Oceanography, 25(1), pp. 18–43. doi: 10.5670/oceanog.2012.02.

Franke, H. D., Gutow, L. and Janke, M. (2007) ‘Flexible habitat selection and interactive habitat segregation in the marine congeners Idotea baltica and Idotea emarginata (Crustacea, Isopoda)’, Marine Biology, 150(5), pp. 929–939. doi: 10.1007/s00227-006-0421-2.

Gagliano, M. and McCormick, M. I. (2007) ‘Compensating in the wild: Is flexible growth the key to early juvenile survival?’, Oikos, 116(1), pp. 111–120. doi: 10.1111/j.2006.0030-1299.15418.x.

Gebruk, A. et al. (2010) ‘Community dynamics over a decadal scale at Logatchev, 14°45’N, Mid-Atlantic Ridge’, Cahiers de Biologie Marine, 51(4), pp. 383–388. doi: 10.21411/CBM.A.FB805CDE.

Gebruk, A. V. et al. (2000) ‘Food sources, behaviour, and distribution of hydrothermal vent shrimps at the Mid-Atlantic Ridge’, Journal of the Marine Biological Association of the United Kingdom, 80(3), pp. 485–499. doi: 10.1017/S0025315400002186.

Geret, F. et al. (2002) ‘Metal bioaccumulation and storage forms in the shrimp, Rimicaris exoculata, from the Rainbow hydrothermal field (Mid-Atlantic Ridge); preliminary approach to the fluid-organism relationship’, Cahiers de Biologie Marine, 43(1), pp. 43–52.

Girard, F. et al. (2020) ‘Currents and topography drive assemblage distribution on an active hydrothermal edifice’, Progress in Oceanography, 187(July), p. 102397. doi: 10.1016/j.pocean.2020.102397.

Gosselin, L. A. and Qian, P. Y. (1997) ‘Juvenile mortality in benthic marine invertebrates’, Marine Ecology Progress Series, 146(1–3), pp. 265–282. doi: 10.3354/meps146265.

Hernández-Ávila, I. et al. (2021) ‘Population structure and reproduction of the alvinocaridid shrimp Rimicaris exoculata on the Mid-Atlantic Ridge: variations between habitats and vent fields’, bioRxiv, pp. 3–5.

Husson, B. et al. (2017) ‘Picturing thermal niches and biomass of hydrothermal vent species’, Deep-Sea Research Part II: Topical Studies in Oceanography, 137, pp. 6–25. doi: 10.1016/j.dsr2.2016.05.028.

Jollivet, D. et al. (2000) ‘Reproductive biology, sexual dimorphism, and population structure of the deep sea hydrothermal vent scale-worm, Branchipolynoe seepensis (Polychaeta:Polynoidae)’, Journal of the Marine Biological Association of the United Kingdom, 80, pp. 55–68. doi: 10.1017/S0025315499001563.

Jones, L. A. and Ricciardi, A. (2014) ‘The influence of pre-settlement and early post-settlement processes on the adult distribution and relative dominance of two invasive mussel species’, Freshwater Biology, 59(5), pp. 1086–1100. doi: 10.1111/fwb.12331.

Kelly, N. E. and Metaxas, A. (2007) ‘Population structure of two deep-sea hydrothermal vent gastropods from the Juan de Fuca Ridge, NE Pacific’, Marine Biology, 153(3), pp. 457–471. doi: 10.1007/s00227-007-0828-4.

Komai, T. and Segonzac, M. (2008) ‘Taxonomic Review of the Hydrothermal Vent Shrimp Genera Rimicaris Williams & Rona and Chorocaris Martin & Hessler (Crustacea: Decapoda: Caridea: Alvinocarididae)’, Journal of Shellfish Research, 27(1), pp. 21–41. doi: 10.2983/0730-8000(2008)27[21:TROTHV]2.0.CO;2.

Lenihan, H. S. et al. (2008) ‘Biotic interactions at hydrothermal vents: Recruitment inhibition by the mussel Bathymodiolus thermophilus’, Deep-Sea Research Part I: Oceanographic Research Papers, 55(12), pp. 1707–1717. doi: 10.1016/j.dsr.2008.07.007.

Levesque, C., Juniper, S. K. and Marcus, J. (2003) ‘Food resource partitioning and competition among alvinellid polychaetes of Juan de Fuca Ridge hydrothermal vents’, Marine Ecology Progress Series, 246(May 2014), pp. 173–182. doi: 10.3354/meps246173.

Macdonald, P. and Du, J. (2012) ‘Mixdist: finite mixture distribution models’, R package version 0.5-4.

Mace, M. M. and Rozas, L. P. (2018) ‘Fish Predation on Juvenile Penaeid Shrimp: Examining Relative Predator Impact and Size-Selective Predation’, Estuaries and Coasts, 41(7), pp. 2128–2134. doi: 10.1007/s12237-018-0409-4.

Marsh, L. et al. (2015) ‘In hot and cold water: Differential life-history traits are key to success in contrasting thermal deep-sea environments’, Journal of Animal Ecology, 84(4), pp. 898– 913. doi: 10.1111/1365-2656.12337.

Marticorena, J. et al. (2020) ‘Contrasting reproductive biology of two hydrothermal gastropods from the Mid-Atlantic Ridge:implications for resilience of vent communities’, Marine Biology, 167(109), pp. 1–19. doi: 10.1007/s00227-020-03721-x.

Matabos, M. and Thiebaut, E. (2010) ‘Reproductive biology of three hydrothermal vent peltospirid gastropods (Nodopelta heminoda, N. subnoda and Peltospira operculata) associated with Pompeii worms on the East Pacific Rise’, Journal of Molluscan Studies, 76(3), pp. 257–266. doi: 10.1093/mollus/eyq008.

Matias, M. G. et al. (2012) ‘Increasing density of rare species of intertidal gastropods: Tests of competitive ability compared with common species’, Marine Ecology Progress Series, 453, pp. 107–116. doi: 10.3354/meps09639.

Meekan, M. G. et al. (2006) ‘Bigger is better: Size-selective mortality throughout the life history of a fast-growing clupeid, Spratelloides gracilis’, Marine Ecology Progress Series, 317(Anderson 1988), pp. 237–244. doi: 10.3354/meps317237.

Menge, B. A. (1991) ‘Relative importance of recruitment and other causes of variation in rocky intertidal community structure’, Journal of Experimental Marine Biology and Ecology, 146, pp. 69–100.

Methou, P. (2019) Lifecycles of two hydrothermal vent shrimps from the Mid-Atlantic Ridge: Rimicaris exoculata and Rimicaris chacei.

Methou, P. et al. (2020) ‘Integrative taxonomy revisits the ontogeny and trophic niches of Rimicaris vent shrimps’, Royal Society Open Science, 7(7), p. 200837. doi: 10.1098/rsos.200837.

Münzbergová, Z. (2013) ‘Comparative demography of two co-occurring Linum species with different distribution patterns’, Plant Biology, 15(6), pp. 963–970. doi: 10.1111/plb.12007.

Nakamura, K. and Takai, K. (2014) ‘Theoretical constraints of physical and chemical properties of hydrothermal fluids on variations in chemolithotrophic microbial communities in seafloor hydrothermal systems’, Progress in Earth and Planetary Science, 1(1), pp. 1–24. doi: 10.1186/2197-4284-1-5.

Nye, V., Copley, J. T. P. and Tyler, P. A. (2013) ‘Spatial Variation in the Population Structure and Reproductive Biology of Rimicaris hybisae (Caridea: Alvinocarididae) at Hydrothermal Vents on the Mid-Cayman Spreading Centre’, PLoS ONE, 8(3). doi: 10.1371/journal.pone.0060319.

Plouviez, S. et al. (2015) ‘Characterization of vent fauna at the mid-cayman spreading center’, Deep-Sea Research Part I: Oceanographic Research Papers, 97, pp. 124–133. doi: 10.1016/j.dsr.2014.11.011.

Ponsard, J. et al. (2013) ‘Inorganic carbon fixation by chemosynthetic ectosymbionts and nutritional transfers to the hydrothermal vent host-shrimp Rimicaris exoculata.’, ISME journal, 7(1), pp. 96–109. doi: 10.1038/ismej.2012.87.

R Core Team,. (2020) R: A language and environment for statistical computing. Vienna, Austria. Available at: https://www.r-project.org.

Ravaux, J. et al. (2016) ‘Plasticity and acquisition of the thermal tolerance (upper thermal limit and heat shock response) in the intertidal species Palaemon elegans’, Journal of Experimental Marine Biology and Ecology, 484, pp. 39–45. doi: 10.1016/j.jembe.2016.07.003.

Ravaux, J. et al. (2019) ‘Assessing a species thermal tolerance through a multiparameter approach: the case study of the deep-sea hydrothermal vent shrimp Rimicaris exoculata’, Cell Stress and Chaperones, 24(3), pp. 647–659. doi: 10.1007/s12192-019-01003-0.

Rius, M., Turon, X. and Marshall, D. J. (2009) ‘Non-lethal effects of an invasive species in the marine environment: The importance of early life-history stages’, Oecologia, 159(4), pp. 873–882. doi: 10.1007/s00442-008-1256-y.

Rybakova, E. and Galkin, S. (2015) ‘Hydrothermal assemblages associated with different foundation species on the East Pacific Rise and Mid-Atlantic Ridge, with a special focus on mytilids’, Marine Ecology, 36(S1), pp. 45–61. doi: 10.1111/maec.12262.

Sarradin, P. M. et al. (2007) ‘Dissolved and particulate metals (Fe, Zn, Cu, Cd, Pb) in two habitats from an active hydrothermal field on the EPR at 13°N’, Science of the Total Environment, 392(1), pp. 119–129. doi: 10.1016/j.scitotenv.2007.11.015.

Schmidt, C., Le Bris, N. and Gaill, F. (2008) ‘Interactions of Deep-Sea Vent Invertebrates with Their Environment: The Case of Rimicaris exoculata’, Journal of Shellfish Research, 27(1), pp. 79–90. doi: 10.2983/0730-8000(2008)27[79:IODVIW]2.0.CO;2.

Segonzac, M., de Saint Laurent, M. and Casanova, B. (1993) ‘L’enigme du comportement trophique des crevettes Alvinocarididae des sites hydrothermaux de la dorsale medio-atlantique’, Cahiers de Biologie Marine, 34(4), pp. 535–571. doi: 10.21411/CBM.A.B3683E29.

Shank, T. M., Lutz, R. a and Vrijenhoek, R. C. (1998) ‘Molecular systematics of shrimp (Decapoda: Bresiliidae) from deep-sea hydrothermal vents, I: Enigmatic “small orange” shrimp from the Mid-Atlantic Ridge are juvenile Rimicaris exoculata.’, Molecular Marine Biology and Biotechnology, 7(2), pp. 88–96.

Shillito, B. et al. (2006) ‘Temperature resistance studies on the deep-sea vent shrimp Mirocaris fortunata’, Journal of Experimental Biology, 209(5), pp. 945–955. doi: 10.1242/jeb.02102.

Stookey, L. L. (1970) ‘Ferrozine - a new spectrophotometric reagent for iron’, Analytical Chemistry, 42(7), pp. 779–781. doi: 10.1021/ac60289a016.

Tibshirani, R., Walther, G. and Hastie, T. (2001) ‘Estimating the number of data clusters via the gap statistic’, Journal of the Royal Statistical Society: Series B, pp. 411–423.

Vereshchaka, A. L., Kulagin, D. N. and Lunina, A. A. (2015) ‘Phylogeny and new classification of hydrothermal vent and seep shrimps of the family alvinocarididae (Decapoda)’, PLoS ONE, 10(7), pp. 1–29. doi: 10.1371/journal.pone.0129975.

Vuillemin, R. et al. (2009) ‘CHEMINI: A new in situ CHEmical MINIaturized analyzer’, Deep-Sea Research Part I: Oceanographic Research Papers, 56(8), pp. 1391–1399. doi: 10.1016/j.dsr.2009.02.002.

Wahle, R. and Steneck, R. (1991) ‘Recruitment habitats and nursery grounds of the American lobster Homarus americanus: a demographic bottleneck?’, Marine Ecology Progress Series, 69(3), pp. 231–243. doi: 10.3354/meps069231.

Watanabe, H. and Beedessee, G. (2015) ‘Vent Fauna on the Central Indian Ridge’, in Subseafloor Biosphere Linked to Hydrothermal Systems: TAIGA Concept, pp. 1–666. doi: 10.1007/978-4-431-54865-2.

Werner, E. E. and Gilliam, J. F. (1984) ‘The Ontogenetic niche and species interactions in size-structured populations’, Annual Review of Ecology and Systematics, 15(1), pp. 393–425. doi: 10.1146/annurev.es.15.110184.002141.

Zal, F. et al. (1995) ‘Reproductive biology and population structure of the deep-sea hydrothermal vent worm Paralvinella grasslei (Polychaeta: Alvinellidae) at 13°N on the East Pacific Rise’, Marine Biology, 122(4), pp. 637–648. doi: 10.1007/BF00350685.

Zbinden, M. et al. (2004) ‘Distribution of bacteria and associated minerals in the gill chamber of the vent shrimp Rimicaris exoculata and related biogeochemical processes’, Marine Ecology Progress Series, 284, pp. 237–251. doi: 10.3354/meps284237.

Zbinden, M. and Cambon-Bonavita, M. (2020) ‘Biology and ecology of Rimicaris exoculata, a symbiotic shrimp from deep-sea hydrothermal vents’, Marine Ecology Progress Series, 652, pp. 187–222. doi: 10.3354/meps13467.

