## Supplementary material for "Population structure and environmental niches of *Rimicaris* shrimps from the Mid Atlantic Ridge": suplemental_material.pdf

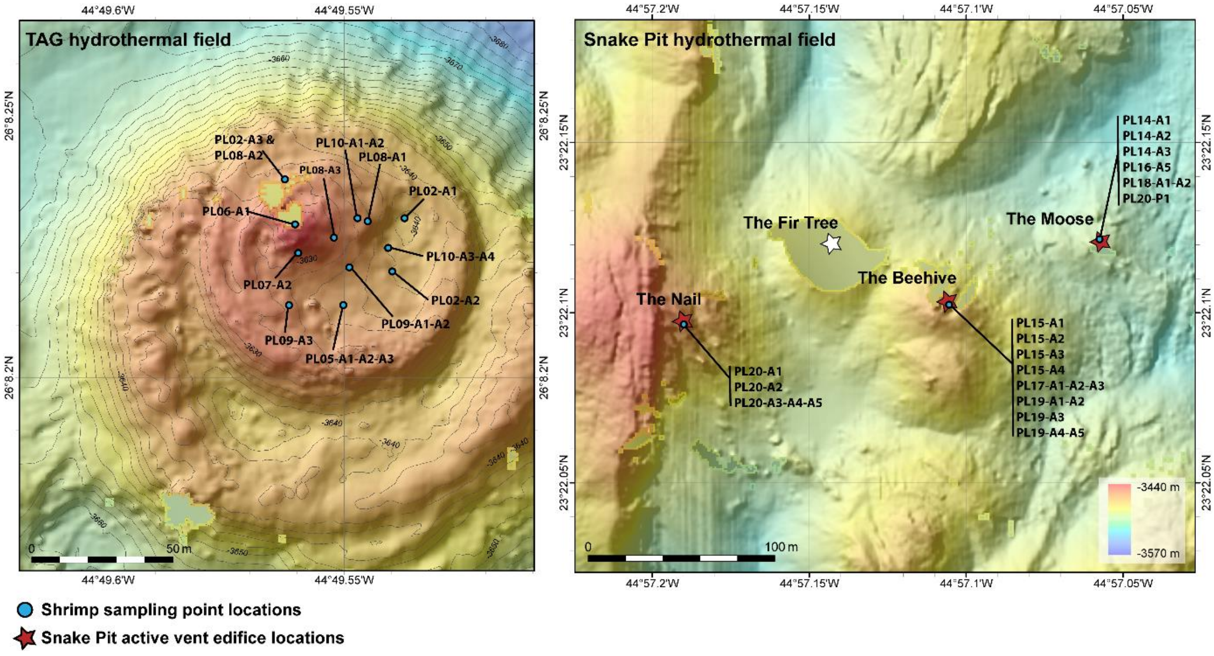


**Figure S1.** Map depicting the different spatially discrete sampling point at the TAG and Snake Pit hydrothermal vent fields

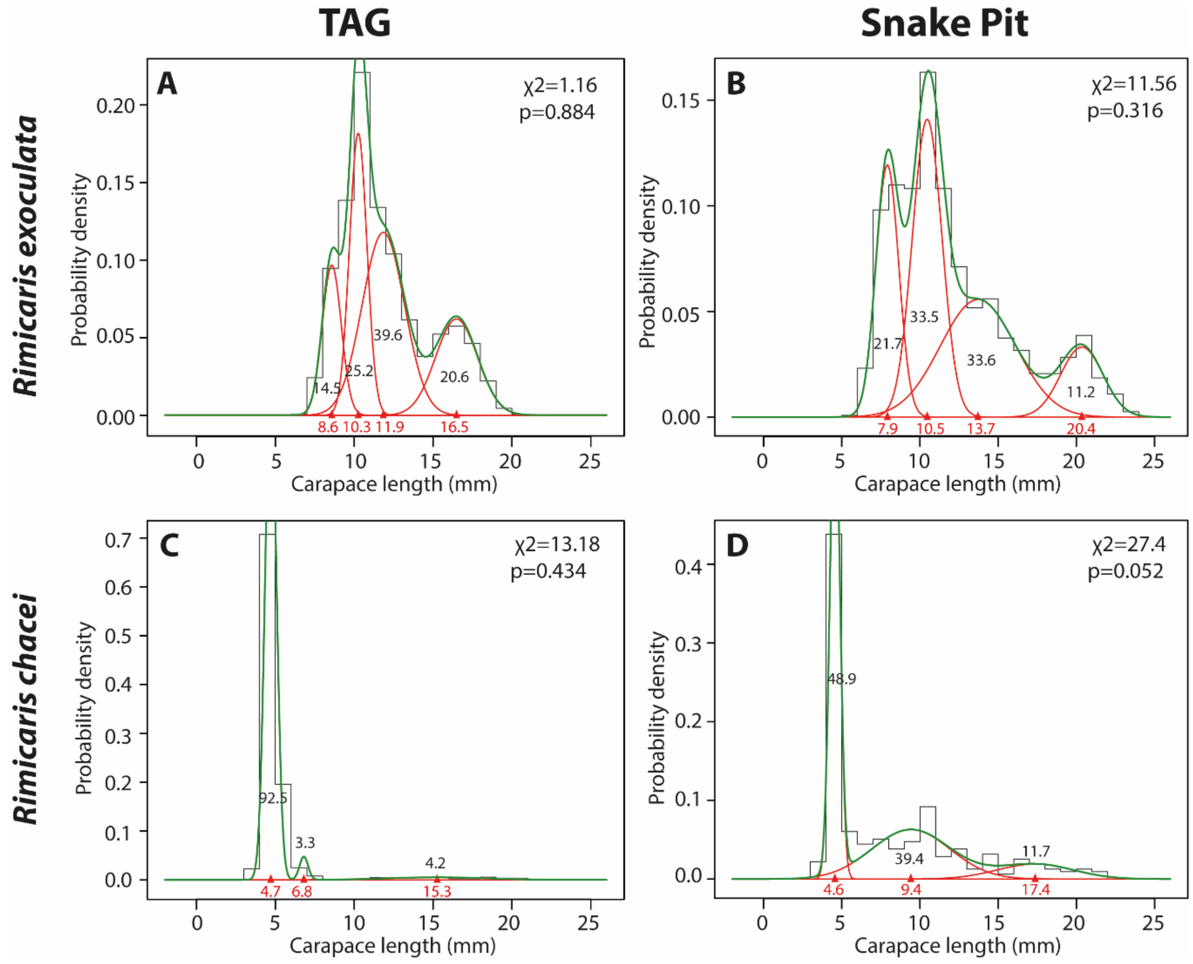


**Figure S2.** Cohort detection through modal decomposition of size-frequency distributions of *Rimicaris* populations. **A.** *R. exoculata* population from TAG. **B.** *R. exoculata* population from Snake Pit. **C.** *R. chacei* population from TAG. **D.** *R. chacei* population from Snake Pit.

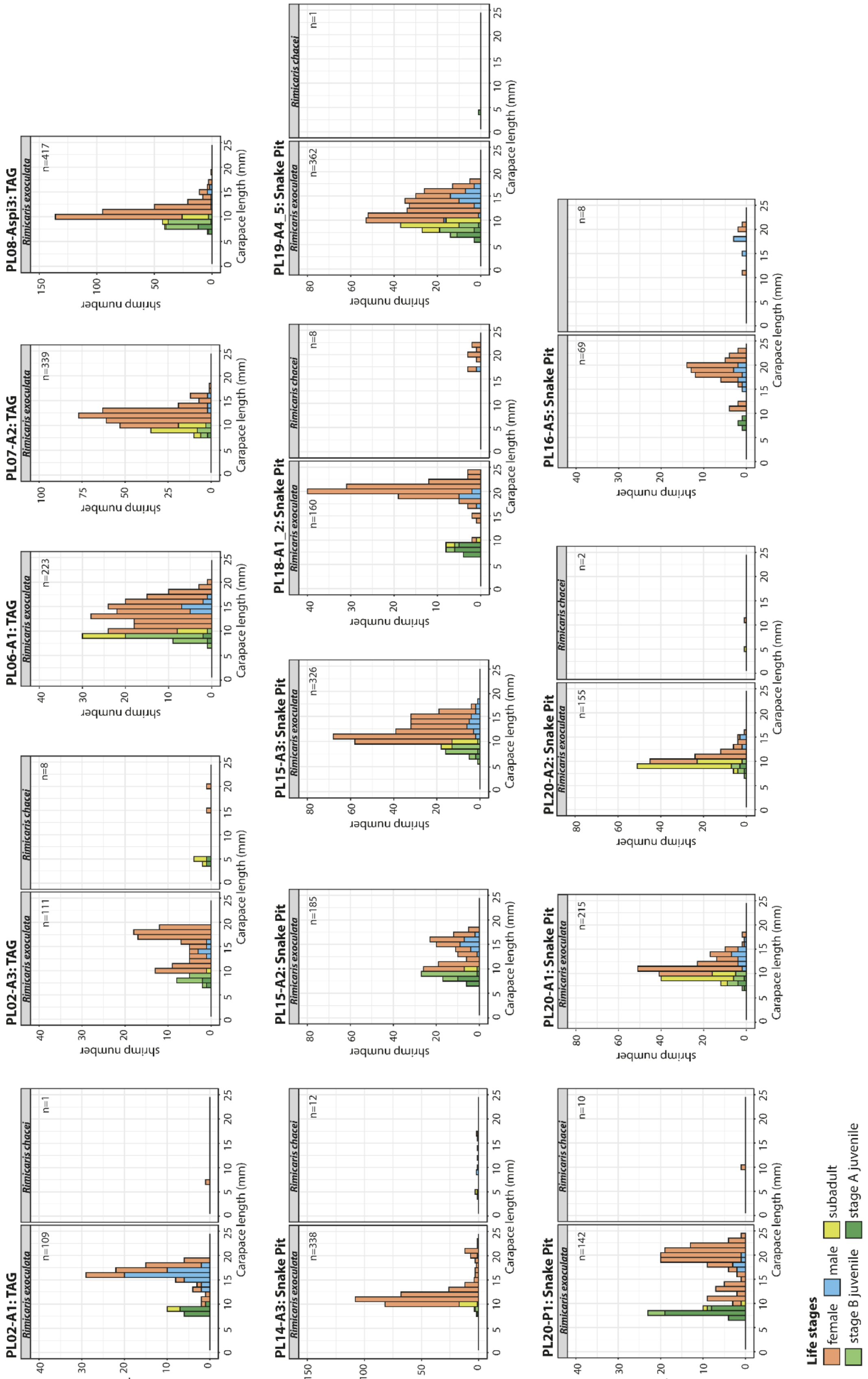


**Figure S3.** Small scale spatial variations in size-frequency distribution between the different dense aggregate assemblages at TAG and Snake Pit.

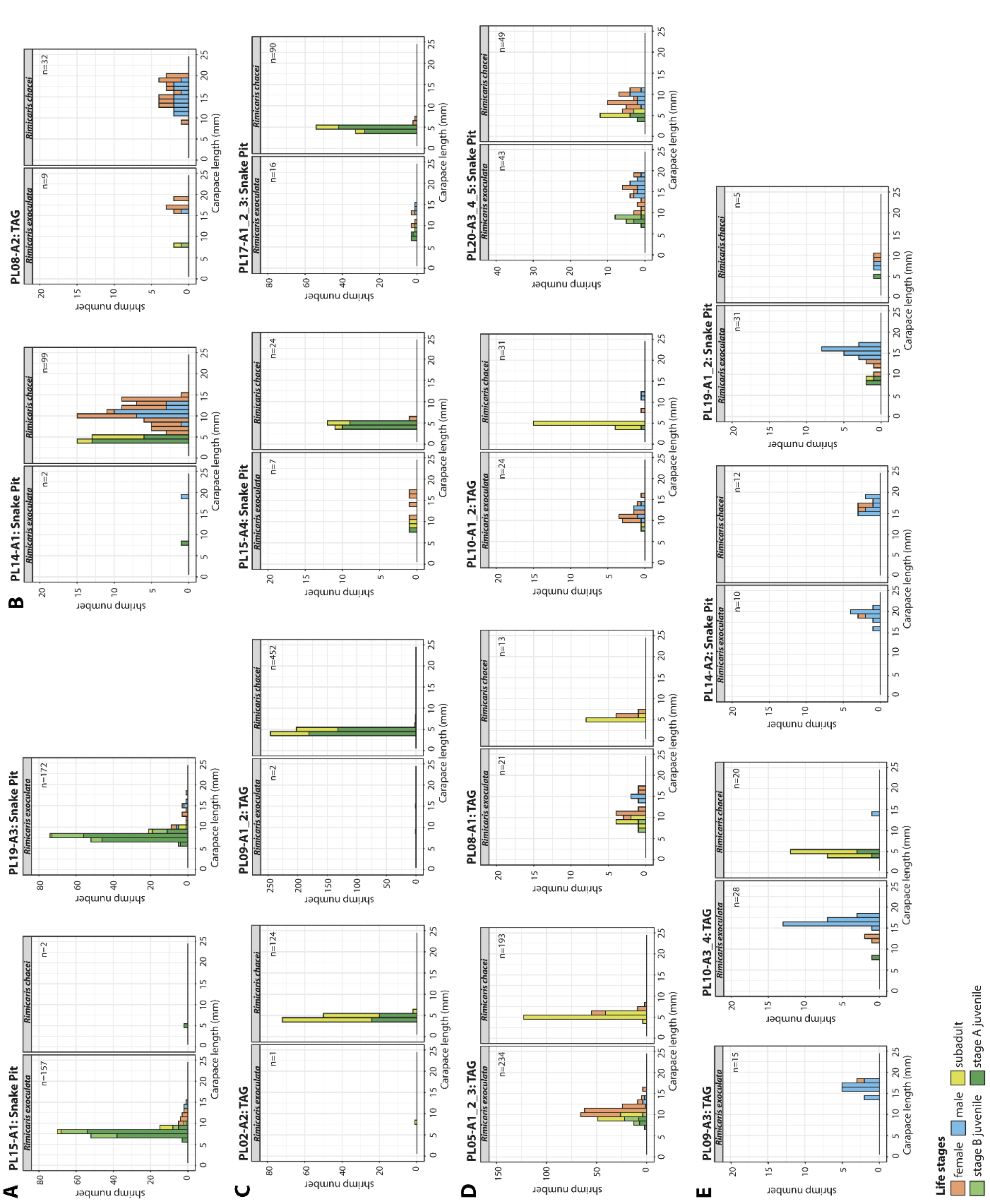


**Figure S4.** Small scale spatial variations in size-frequency distribution between the different assemblage types at TAG and Snake Pit. **A.** Nurseries of R. exoculata **B.** Hidden aggregates of R. chacei adults **C.** Nurseries of R. chacei **D.** Scattered assemblages in inactive habitats **E.** Mixed shrimps assemblages at the periphery of dense aggregates.

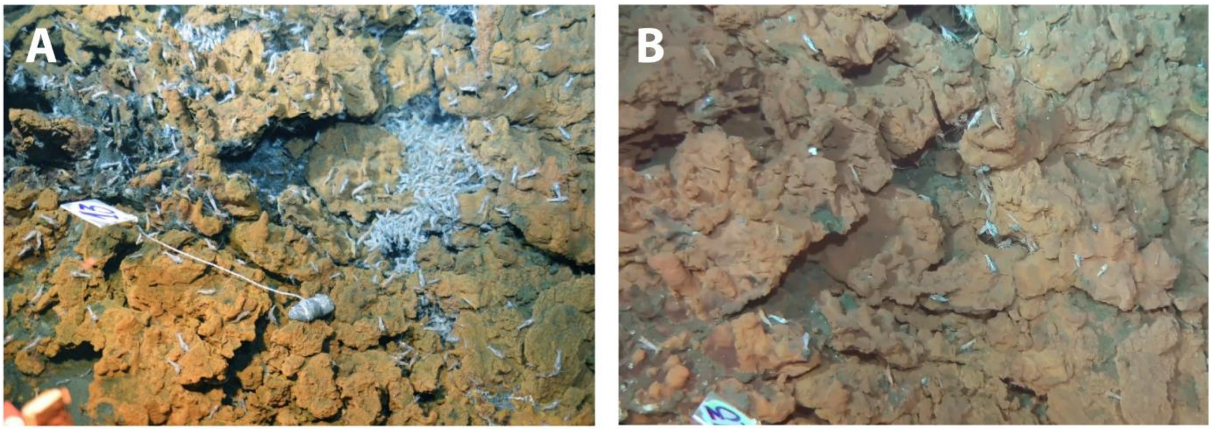


**Figure S5.** Temporal succession of assemblages observed at TAG. **A.** Dense aggregate of *R. exoculata* adults **B.** Hidden aggregate of *R. chacei* adults observed at the same location 10 days later.

| Vent field | Localisation | Assemblage type | Sample ID | *Rimicaris exoculata* | | | | | | | | *Rimicaris chacei* | | | | | | |
|---|---|---|---|---|---|---|---|---|---|---|---|---|---|---|---|---|---|---|
| | | | | F | M | S | JB | JA | Ad | Im | **n** | F | M | S | JA | Ad | Im | **n** |
| TAG | near active fluids | Dense Aggregate | PL02-A1 | 50 | 42 | 3 | 0 | 14 | 92 | 17 | **109** | 1 | 0 | 0 | 0 | 1 | 0 | **1** |
| TAG | near active fluids | Dense Aggregate | PL02-A3 | 89 | 6 | 1 | 12 | 3 | 95 | 16 | **111** | 2 | 0 | 4 | 2 | 2 | 6 | **8** |
| TAG | near active fluids | Dense Aggregate | PL06-A1 | 160 | 15 | 17 | 28 | 3 | 175 | 48 | **223** | 0 | 0 | 0 | 0 | 0 | 0 | **0** |
| TAG | near active fluids | Dense Aggregate | PL07-A2 | 269 | 6 | 47 | 15 | 2 | 275 | 64 | **339** | 0 | 0 | 0 | 0 | 0 | 0 | **0** |
| TAG | near active fluids | Dense Aggregate | PL08-A3 | 294 | 9 | 29 | 69 | 16 | 303 | 114 | **417** | 0 | 0 | 0 | 0 | 0 | 0 | **0** |
| Snake Pit | near active fluids | Dense Aggregate | PL14-A3 | 313 | 1 | 16 | 5 | 1 | 314 | 22 | **336** | 2 | 1 | 0 | 0 | 3 | 0 | **3** |
| Snake Pit | near active fluids | Dense Aggregate | PL15-A2 | 103 | 23 | 6 | 34 | 19 | 126 | 59 | **185** | 0 | 0 | 0 | 0 | 0 | 0 | **0** |
| Snake Pit | near active fluids | Dense Aggregate | PL15-A3 | 247 | 25 | 17 | 34 | 3 | 272 | 54 | **326** | 0 | 0 | 0 | 0 | 0 | 0 | **0** |
| Snake Pit | near active fluids | Dense Aggregate | PL16-A5 | 55 | 10 | 0 | 0 | 4 | 65 | 4 | **69** | 4 | 4 | 0 | 0 | 8 | 0 | **8** |
| Snake Pit | near active fluids | Dense Aggregate | PL18-A1A2 | 113 | 8 | 3 | 3 | 15 | 121 | 21 | **142** | 9 | 1 | 0 | 0 | 10 | 0 | **10** |
| Snake Pit | near active fluids | Dense Aggregate | PL19-A4A5 | 226 | 39 | 51 | 28 | 18 | 265 | 97 | **362** | 0 | 0 | 0 | 1 | 0 | 1 | **1** |
| Snake Pit | near active fluids | Dense Aggregate | PL20-P1 | 115 | 7 | 2 | 5 | 31 | 122 | 38 | **160** | 1 | 0 | 0 | 0 | 1 | 0 | **1** |
| Snake Pit | near active fluids | Dense Aggregate | PL20-A1 | 122 | 23 | 48 | 16 | 6 | 145 | 70 | **215** | 0 | 0 | 0 | 0 | 0 | 0 | **0** |
| Snake Pit | near active fluids | Dense Aggregate | PL20-A2 | 69 | 5 | 67 | 10 | 4 | 74 | 81 | **155** | 1 | 0 | 1 | 0 | 1 | 1 | **2** |
| Snake Pit | near diffuse fluids | *R. exoculata* Nursery | PL15-A1 | 12 | 3 | 10 | 32 | 100 | 15 | 142 | **157** | 0 | 0 | 0 | 2 | 0 | 2 | **2** |
| Snake Pit | near diffuse fluids | *R. exoculata* Nursery | PL19-A3 | 11 | 4 | 7 | 33 | 117 | 15 | 157 | **172** | 0 | 0 | 0 | 0 | 0 | 0 | **0** |
| TAG | near diffuse fluids | Hidden Aggregate | PL08-A2 | 6 | 1 | 1 | 1 | 0 | 7 | 2 | **9** | 16 | 16 | 0 | 0 | 32 | 0 | **32** |
| Snake Pit | near diffuse fluids | Hidden Aggregate | PL14-A1 | 0 | 1 | 0 | 0 | 1 | 1 | 1 | **2** | 47 | 24 | 9 | 19 | 71 | 28 | **99** |
| TAG | near diffuse fluids | *R. chacei* Nursery | PL02-A2 | 0 | 0 | 1 | 0 | 0 | 0 | 1 | **1** | 0 | 0 | 80 | 44 | 0 | 124 | **124** |
| TAG | near diffuse fluids | *R. chacei* Nursery | PL09-A1A2 | 0 | 1 | 0 | 0 | 1 | 1 | 1 | **2** | 4 | 0 | 135 | 313 | 4 | 448 | **452** |
| Snake Pit | near diffuse fluids | *R. chacei* Nursery | PL15-A4 | 4 | 0 | 2 | 0 | 1 | 4 | 3 | **7** | 1 | 0 | 4 | 19 | 1 | 23 | **24** |
| Snake Pit | near diffuse fluids | *R. chacei* Nursery | PL17-A1A2A3 | 5 | 3 | 0 | 3 | 5 | 8 | 8 | **16** | 3 | 0 | 17 | 70 | 3 | 87 | **90** |
| TAG | near diffuse fluids | Vicinity of D. Aggregates | PL05-A1A2A3 | 137 | 7 | 49 | 24 | 17 | 144 | 90 | **234** | 25 | 0 | 168 | 0 | 25 | 168 | **193** |
| TAG | near diffuse fluids | Vicinity of D. Aggregates | PL08-A1 | 9 | 3 | 6 | 3 | 0 | 12 | 9 | **21** | 4 | 0 | 9 | 0 | 4 | 9 | **13** |
| TAG | near diffuse fluids | Vicinity of D. Aggregates | PL10-A1A2 | 15 | 7 | 1 | 1 | 0 | 22 | 2 | **24** | 1 | 2 | 37 | 1 | 3 | 38 | **41** |
| Snake Pit | near diffuse fluids | | PL20-A3A4A5 | 6 | 5 | 0 | 5 | 4 | 11 | 9 | **20** | 4 | 5 | 8 | 6 | 9 | 14 | **23** |
| TAG | far from fluid activity | Scattered individuals | PL09-A3 | 1 | 14 | 0 | 0 | 0 | 15 | 0 | **15** | 0 | 0 | 0 | 0 | 0 | 0 | **0** |
| TAG | far from fluid activity | Scattered individuals | PL10-A3A4 | 3 | 24 | 0 | 0 | 1 | 27 | 1 | **28** | 0 | 1 | 15 | 4 | 1 | 19 | **20** |
| Snake Pit | far from fluid activity | Scattered individuals | PL14-A2 | 2 | 9 | 1 | 0 | 0 | 11 | 1 | **12** | 3 | 13 | 2 | 2 | 16 | 4 | **20** |
| Snake Pit | far from fluid activity | Scattered individuals | PL19-A1A2 | 4 | 19 | 1 | 0 | 3 | 23 | 4 | **27** | 2 | 2 | 0 | 1 | 4 | 1 | **5** |

**Table S1.** *Rimicaris* life stage composition of each assemblages.

| Vent field | Localisation | Assemblage type | Sample ID | Specie ratio | | | Juvenile ratio | | | Sex ratio | | | Juvenile ratio | | | Sex ratio | | |
|---|---|---|---|---|---|---|---|---|---|---|---|---|---|---|---|---|---|---|
| | | | | TRch:TRex | χ2 (1 df) | Significance | TI:TA | χ2 (1 df) | Significance | TM:TF | χ2 (1 df) | Significance | TI:TA | χ2 (1 df) | Significance | TM:TF | χ2 (1 df) | Significance |
| TAG | near active fluids | Dense Aggregate | PL02-A1 | 0.009:1 | 208.2 | *** | 0.18:1 | 100.5 | *** | 0.84:1 | 0.69 | NS | - | - | - | - | - | - |
| TAG | near active fluids | Dense Aggregate | PL02-A3 | 0.07:1 | 174.9 | *** | 0.17:1 | 109.6 | *** | 0.07:1 | 72.5 | *** | - | - | - | - | - | - |
| TAG | near active fluids | Dense Aggregate | PL06-A1 | 0.0:1 | - | - | 0.27:1 | 142.4 | *** | 0.09:1 | 120.1 | *** | - | - | - | - | - | - |
| TAG | near active fluids | Dense Aggregate | PL07-A2 | 0.0:1 | - | - | 0.23:1 | 260.2 | *** | 0.02:1 | 251.5 | *** | - | - | - | - | - | - |
| TAG | near active fluids | Dense Aggregate | PL08-A3 | 0.0:1 | - | - | 0.38:1 | 169.5 | *** | 0.03:1 | 268.1 | *** | - | - | - | - | - | - |
| Snake Pit | near active fluids | Dense Aggregate | PL14-A3 | 0.009:1 | 650.3 | *** | 0.07:1 | 504.1 | *** | 0.003:1 | 310 | *** | - | - | - | - | - | - |
| Snake Pit | near active fluids | Dense Aggregate | PL15-A2 | 0.0:1 | - | - | 0.47:1 | 47.1 | *** | 0.22:1 | 50.8 | *** | - | - | - | - | - | - |
| Snake Pit | near active fluids | Dense Aggregate | PL15-A3 | 0.0:1 | - | - | 0.20:1 | 288.9 | *** | 0.10:1 | 181.2 | *** | - | - | - | - | - | - |
| Snake Pit | near active fluids | Dense Aggregate | PL16-A5 | 0.12:1 | 93.5 | *** | 0.06:1 | 104.4 | *** | 0.18:1 | 31.2 | *** | - | - | - | - | - | - |
| Snake Pit | near active fluids | Dense Aggregate | PL18-A1A2 | 0.07:1 | 225.8 | *** | 0.17:1 | 138.1 | *** | 0.07:1 | 95.6 | *** | 0.0:1 | 16.2 | *** | 0.11:1 | 6.4 | * |
| Snake Pit | near active fluids | Dense Aggregate | PL19-A4A5 | 0.003:1 | 714.1 | *** | 0.37:1 | 154.1 | *** | 0.17:1 | 132 | *** | - | - | - | - | - | - |
| Snake Pit | near active fluids | Dense Aggregate | PL20-P1 | 0.006:1 | 310.1 | *** | 0.31:1 | 86.1 | *** | 0.06:1 | 91.1 | *** | - | - | - | - | - | - |
| Snake Pit | near active fluids | Dense Aggregate | PL20-A1 | 0.0:1 | - | - | 0.48:1 | 50.9 | *** | 0.19:1 | 67.6 | *** | - | - | - | - | - | - |
| Snake Pit | near active fluids | Dense Aggregate | PL20-A2 | 0.01:1 | 294.3 | *** | 1.09:1 | 0.5 | NS | 0.07:1 | 55.4 | *** | - | - | - | - | - | - |
| Snake Pit | near diffuse fluids | *R. exoculata* Nursery | PL15-A1 | 0.01:1 | 298.3 | *** | 9.47:1 | 202.2 | *** | 0.25:1 | 5.4 | * | - | - | - | - | - | - |
| Snake Pit | near diffuse fluids | *R. exoculata* Nursery | PL19-A3 | 0.0:1 | - | - | 10.47:1 | 231.2 | *** | 0.36:1 | 3.2 | NS | - | - | - | - | - | - |
| TAG | near diffuse fluids | Hidden Aggregate | PL08-A2 | 3.56:1 | 23.6 | *** | - | - | - | - | - | - | 0.0:1 | 60.1 | *** | 1.0:1 | 0 | NS |
| Snake Pit | near diffuse fluids | Hidden Aggregate | PL14-A1 | 49.5:1 | 182.5 | *** | - | - | - | - | - | - | 0.39:1 | 35.6 | *** | 0.51:1 | 7.4 | ** |
| TAG | near diffuse fluids | *R. chacei* Nursery | PL02-A2 | 124.0:1 | 238.1 | *** | - | - | - | - | - | - | 124.0:0 | 244 | *** | - | - | - |
| TAG | near diffuse fluids | *R. chacei* Nursery | PL09-A1A2 | 226.0:1 | 888.1 | *** | - | - | - | - | - | - | 112.0:1 | 868.4 | *** | - | - | - |
| Snake Pit | near diffuse fluids | *R. chacei* Nursery | PL15-A4 | 3.43:1 | 16.5 | *** | - | - | - | - | - | - | 23.0:1 | 36.8 | *** | - | - | - |
| Snake Pit | near diffuse fluids | *R. chacei* Nursery | PL17-A1A2A3 | 5.63:1 | 100.6 | *** | 1.0:1 | 0.0 | NS | - | - | - | 29.0:1 | 153.1 | *** | - | - | - |
| TAG | near diffuse fluids | Vicinity of D. Aggregates | PL05-A1A2A3 | 0.82:1 | 7.5 | * | 0.63:1 | 24 | *** | 0.05:1 | 117.4 | *** | 6.72:1 | 208.9 | *** | 0.0:1 | 25 | *** |
| TAG | near diffuse fluids | Vicinity of D. Aggregates | PL08-A1 | 0.62:1 | 2.9 | NS | 0.75:1 | 0.4 | NS | 0.33:1 | 3 | NS | 2.25:1 | 2.5 | NS | - | - | - |
| TAG | near diffuse fluids | Vicinity of D. Aggregates | PL10-A1A2 | 1.71:1 | 7.9 | * | 0.09:1 | 30.1 | *** | 0.47:1 | 2.9 | NS | 12.67:1 | 56.4 | *** | - | - | - |
| TAG | far from fluid activity | Scattered individuals | PL09-A3 | 0.0:1 | - | - | 0.0:1 | 26.1 | *** | 14.0:1 | 11.3 | *** | - | - | - | - | - | - |
| TAG | far from fluid activity | Scattered individuals | PL10-A3A4 | 0.7:1 | 2 | NS | 0.04:1 | 44.6 | *** | 8.0:1 | 16.3 | *** | 19.0:1 | 28.9 | *** | - | - | - |
| Snake Pit | far from fluid activity | Scattered individuals | PL14-A2 | 1.7:1 | 3.6 | NS | 0.0:1 | 16.2 | *** | 4.5:1 | 4.45 | * | 0.25:1 | 12.1 | *** | 0.23:1 | 6.25 | * |
| Snake Pit | far from fluid activity | Scattered individuals | PL19-A1A2 | 0.19:1 | 27.6 | *** | 0.17:1 | 24 | *** | 4.75:1 | 9.8 | ** | - | - | - | - | - | - |
| Snake Pit | near diffuse fluids | - | PL20-A3A4A5 | 1.15:1 | 0.2 | NS | 0.82:1 | 0.1 | NS | 0.83:1 | 0.1 | NS | 1.56:1 | 1.4 | NS | 1.25:1 | 0.11 | NS |

**Table S2.** Spatial variation in the sex ratio, juvenile ratio and the proportions of the different *Rimicaris* species within the different assemblages.

| Vent Field | Assemblage Type | Sample ID | Measurement time | Avg.T (°C) | Std.T (°C) | Min.T (°C) | Max.T (°C) | pH | $H_2S$ (μM) | $Fe^{2+}$ (μM) | $Fe_{total}$ (μM) |
|---|---|---|---|---|---|---|---|---|---|---|---|
| TAG | Dense Aggregate | PL02_A1 | 6min 44s | 8.5 | 4.8 | 3.7 | 20.4 | - | - | - | - |
| TAG | Dense Aggregate | PL06_A1 | 5min 13s | 20.1 | 4.6 | 11.0 | 33.0 | 6.86 | < 1.0 | 239.0 | 315.9 |
| TAG | Dense Aggregate | PL08_A3 | 5min 3s | 9.4 | 4.5 | 5.0 | 19.8 | 7.9 | 3.1 | 30.7 | 129.4 |
| TAG | Hidden Aggregate | PL08_A2 | 5min 10s | 4.9 | 0.7 | 4.5 | 6.9 | 7.7 | 3.7 | 15.3 | 33.1 |
| TAG | Periphery | PL10_A3_4 | 5min 12s | 3.4 | 0.3 | 3.1 | 4.1 | 7.7 | < 1.0 | 10.8 | 179.4 |
| TAG | *R. chacei* nursery | PL02_A2 | 4 min 56s | 4.8 | 0.3 | 4.2 | 5.4 | - | - | - | - |
| TAG | *R. chacei* nursery | PL09_A1_2 | 5min 17s | 3.8 | 0.5 | 3.3 | 5.3 | 7.9 | < 1.0 | n.d. | 9.9 |
| TAG | Vicinity of Aggregates | PL10_A1_2 | 4min 4s | 8.2 | 3.9 | 4.2 | 20.7 | 7.8 | < 1.0 | 47.6 | 75.7 |
| Snake Pit | Dense Aggregate | PL15_A2 | 4min 56s | 7.7 | 0.8 | 6.7 | 10.1 | 7.41 | 1.3 | 0.0 | 22.9 |
| Snake Pit | Dense Aggregate | PL15_A3 | 5min 21s | 20.3 | 5.9 | 11.4 | 38.1 | 7.08 | 8.5 | 11.2 | 34.0 |
| Snake Pit | Dense Aggregate | PL16_A5 | 5min 1s | 6.0 | 1.5 | 3.9 | 8.4 | 7.8 | < 1.0 | 6.2 | 3.0 |
| Snake Pit | Dense Aggregate | PL18_A1_2 | 5min 32s | 9.7 | 2.1 | 5.3 | 13.1 | 7.4 | 1.7 | n.d. | 16.7 |
| Snake Pit | Dense Aggregate | PL19_A4_5 | 6min 14s | 7.0 | 1.3 | 4.9 | 9.5 | 7.36 | 5.7 | 0.0 | 28.8 |
| Snake Pit | Dense Aggregate | PL20_A1 | 5min | 22.5 | 6.9 | 12.1 | 30.7 | - | - | - | - |
| Snake Pit | Dense Aggregate | PL20_A2 | 5min 39s | 14.5 | 1.5 | 8.1 | 19.0 | - | - | - | - |
| Snake Pit | Periphery | PL19_A1_2 | 4min 51s | 3.6 | 0.8 | 3.3 | 6.1 | 7.9 | < 1.0 | 0.2 | < 1.0 |
| Snake Pit | *R. chacei* nursery | PL15_A4 | 3min 1s | 3.9 | 0.6 | 3.4 | 6.8 | - | - | - | - |
| Snake Pit | | PL20_A3_4_5 | 5min 20s | 4.4 | 0.5 | 4.2 | 5.9 | - | - | - | - |
| Snake Pit | *R. exoculata* nursery | PL15_A1 | 5min 4s | 3.4 | 0.5 | 3.2 | 4.9 | 7.8 | < 1.0 | 0.0 | 4.6 |
| Snake Pit | *R. exoculata* nursery | PL19_A3 | 4min 59s | 4.8 | 0.6 | 3.9 | 6.6 | 7.9 | < 1.0 | 0.0 | < 1.0 |

**Table S3.** Temperature and chemistry measurements in each assemblages.

| Species | Vent field | Adults | | | | Females | | | | Male | | | | Subadult | | | | Stage B juveniles | | | | Stage A juveniles | | | |
|---|---|---|---|---|---|---|---|---|---|---|---|---|---|---|---|---|---|---|---|---|---|---|---|---|---|
| | | mean | sd | max | min | mean | sd | max | min | mean | sd | max | min | mean | sd | max | min | mean | sd | max | min | mean | sd | max | min |
| *Rimicaris exoculata* | TAG | 12.9 | 2.6 | 20.1 | 10 | 12.6 | 2.5 | 20.1 | 10 | 15.5 | 1.6 | 18.2 | 10.5 | 9.4 | 0.4 | 9.9 | 7.7 | 8.8 | 0.6 | 10 | 7.2 | 8.3 | 0.6 | 9.7 | 7 |
| | Snake Pit | 14 | 3.6 | 24.4 | 10 | 13.8 | 3.7 | 24.4 | 10 | 15.5 | 2.2 | 20.6 | 10.5 | 9.4 | 0.5 | 9.9 | 7.6 | 8.4 | 0.8 | 10.2 | 6.2 | 7.8 | 0.7 | 9.5 | 5.7 |
| *Rimicaris chacei* | TAG | 11.1 | 5 | 20.5 | 6 | 10 | 5.1 | 20.5 | 6 | 14.3 | 2.6 | 19.1 | 10.9 | 4.9 | 0.5 | 5.9 | 4 | - | - | - | - | 4.5 | 0.3 | 5.4 | 3.8 |
| | Snake Pit | 11.7 | 4.0 | 21.8 | 6.2 | 11.4 | 4.3 | 21.8 | 6.2 | 12.1 | 3.3 | 19 | 7 | 4.9 | 0.4 | 5.9 | 4.1 | - | - | - | - | 4.6 | 0.3 | 5.4 | 3.7 |

**Table S4.** Body size (CL) of the different life stages of *Rimicaris exoculata* and *Rimicaris chacei* from TAG and Snake Pit.
